# The role of memory in non-genetic inheritance and its impact on cancer treatment resistance

**DOI:** 10.1101/2021.02.22.431869

**Authors:** Tyler Cassidy, Daniel Nichol, Mark Robertson-Tessi, Morgan Craig, Alexander R.A. Anderson

## Abstract

Intra-tumour heterogeneity is a leading cause of treatment failure and disease progression in cancer. While genetic mutations have long been accepted as a primary mechanism of generating this heterogeneity, the role of phenotypic plasticity is becoming increasingly apparent as a driver of intra-tumour heterogeneity. Consequently, understanding the role of plasticity in treatment resistance and failure is a key component of improving cancer therapy. We develop a mathematical model of stochastic phenotype switching that tracks the evolution of drug-sensitive and drug-tolerant subpopulations to clarify the role of phenotype switching on population growth rates and tumour persistence. By including cytotoxic therapy in the model, we show that, depending on the strategy of the drug-tolerant subpopulation, stochastic phenotype switching can lead to either transient or permanent drug resistance. We study the role of phenotypic heterogeneity in a drug-resistant, genetically homogeneous population of non-small cell lung cancer cells to derive a rational treatment schedule that drives population extinction and avoids competitive release of the drug-tolerant sub-population. This model-informed therapeutic schedule results in increased treatment efficacy when compared against periodic therapy, and, most importantly, sustained tumour decay without the development of resistance.

**Author summary:** We propose a simple mathematical model to understand the role of phenotypic plasticity and non-genetic inheritance in driving therapy resistance in cancer. We identify the role of non-genetic inheritance on treatment resistance and use a variety of analytical and numerical techniques to understand the impact of phenotypic plasticity on population fitness and dynamics. We further use our model to study the role of phenotypic heterogeneity in therapeutic resistance in a genetically identical non-small cell lung cancer population. Finally, we combine analytical perspectives and techniques from the theory of structured populations, renewal equations and infinite dimensional dynamical systems to derive a model informed therapeutic strategy that both drives tumour eradication while avoiding competitive release of a drug-tolerant subpopulation. These results exemplify the potential of using mathematical techniques to identify therapeutic strategies to guide the evolution of a heterogeneous tumour.

## Introduction

Intra-tumour heterogeneity is a leading driver of cancer treatment failure [1–3]. The genetic instability and high proliferative capacity typical of cancer cells induces a genetically heterogenous population in which resistance-conferring mutations can arise and expand during the selective pressure of therapy. This evolutionary process leads to the eventual failure of treatment and the outgrowth of a refractory tumour [3–9]. However, it is increasingly understood that genetic aberrations are not the sole mechanism through which drug-resistant phenotypes can arise. Rather, adaptive phenotypic change can arise without an associated genetic mutation. Such phenotypic heterogeneity has been extensively studied as a possible mechanism of treatment resistance [7, 8, 10–14]. For example, chemotherapy has been shown to induce a transient drug-tolerant phenotype in breast cancer cell lines such that re-sensitisation occurs following cessation of therapy [15,16], an example of phenotypic plasticity [17]. Whilst this heterogeneity arises in response to environmental change, non-genetic variation in phenotypes can also arise in unchanged environments, indicating the presence of stochastic phenotype switching, termed bet-hedging [13,14]. Bet-hedging induces phenotypic diversity that can help protect a population from extinction following catastrophic environmental changes such as cytotoxic therapy [18–21].

In recent years, evolutionarily-informed cancer therapy regimens have arisen as a potential strategy to delay the emergence of drug resistance. *Adaptive therapies* exploit competition between clonal populations by incorporating periods without therapy wherein resistant subclones, which are often assumed to have a fitness cost in the absence of treatment, can be outcompeted by drug-sensitive clones [5, 22–24]. The treatment is applied and removed based on one or more biomarkers of disease, typically proxy measurements for tumour burden. The theory underlying cancer adaptive therapy is primarily based on competition dynamics between tumour subclones that are not plastic, for example clones arising from mutations. It is presently unclear whether adaptive therapies will prove as effective in mitigating resistance driven by non-genetic mechanisms that change on a faster timescale than mutational rates, or whether better understanding of such non-genetic drivers of resistance could help in the design of more effective evolutionary therapies. Here, we address this question and study the impact of bet-hedging strategies on the development of treatment resistance by developing a simple and qualitative mathematical model.

Mathematical models have been used extensively to understand the development of resistance to anti-cancer therapies. A number of authors have considered how phenotypic variation arises [25,26] as well as the effects of phenotypic heterogeneity (see [11] and references therein). These models rely, in large part, on the use of structured equations which bridge cellular dynamics and population level heterogeneity by explicitly considering the cellular phenotype. Often formulated as partial differential equations (PDEs) structured in phenotypic space, these models conceptualise continuously varying cellular phenotypes [27–30]. These PDE models often include non-local terms to incorporate interactions between cells of different phenotypes, and solutions of these models typically describe the density of cells in phenotype space. Consequently, these structured mathematical models are well-suited to study phenotypic evolution in dynamic environments.

We are particularly interested in the role of stochastic phenotype switching on the development of resistance to anti-cancer therapies. It has previously been shown that stochastic switching between quiescence and proliferation in mammalian cells is biased by the inheritance of mitogen and p53 signalling factors at cell division [31]. The concentration of these factors was shown to be dependent on the life history of the parental cell and are thus representative of non-genetic ‘memory’ in phenotypic switching [31]. Theoretical studies of chemical reaction networks have demonstrated that simple combinations of catalytic and autocatalytic reactions can produce such bistable switches, with different network structures inducing different convergence and stability behaviour [32,33]. Coupled with the inheritence and subsequent decay of intracellular signalling factors, these bistable switches can govern a diverse range of memory-driven switching regimes. Here, we investigate the role of phenotypic memory in intracellular inheritance in driving the emergence of a resistant phenotype during therapy. To this end, we use the cellular age, rather than phenotypic state, to structure our model. In this framework, the cellular age acts as a cipher for a number of epigenetic factors that vary throughout the cell’s life, such as protein accumulation/dilution, cell size, adaptation to environmental stresses, etc. In the model, we use the age of the parent cell to determine the probability that daughter cells will inherit the parental phenotype: cells that reproduce soon after their birth are more likely to bequeath their phenotype to their offspring. This mapping from cellular age to switching probability generalises the role of the bistable switch mechanism that governs phenotypic differentiation, as well any age-driven changes to its behaviour.

The canonical example of the stochastic nature of phenotypic inheritance is the existence of persister cells resulting from stochastic phenotypic switches in *Escherichia coli* populations [14, 20, 34,35]. Comparatively rare in a population in stable exponential growth, the proportion of persister cells increases as the population of *E.coli* cells competes for limited resources [20, 34]. Accordingly, we study the role of growth phase on population composition. Specifically, we demonstrate that populations in exponential and stationary growth stages respond differently to environmental changes. For example, we show that changes in the relative fitness between two phenotypically distinct populations has a drastically different impact on a population in exponential growth compared against the same change during the stationary growth phase. Further, we study the role of phenotypic cooperation on population growth in nutrient-rich environments, and show that this cooperation can hasten population growth when compared against purely Malthusian growth.

We also study the establishment of a resistant population during cytotoxic treatment. Through numerical simulation, we demonstrate that different phenotype switching strategies result in either transient resistance [15,16], or permanent resistance due to the establishment of a dominant, resistant population [36, 37]. We then investigate alternative treatment scheduling options to delay or avoid the establishment of resistance by preserving a drug-sensitive population. This scheduling, inspired by adaptive therapy [5,22], is then shown to outperform periodic or maximally tolerable dosing strategies in a result that is robust to parameter changes. Applying our model to *in vitro* growth data from genetically homogeneous non-small cell lung cancer (NSCLC) populations, we study the effect of phenotypic switching in resistance to anti-cancer drugs. We use two different measurements of fitness under treatment to derive a model-informed therapeutic schedule that balances the desire to drive tumour extinction with the need to avoid competitive release of a drug-tolerant population and the resulting therapy resistance. We show that this treatment schedule leads to long term tumour decay and significantly outperforms metronomic dosing. In the interest of clarity, we present the full analytic results in the Supplemental Information.

## Results

### Phenotypic switching model

Our primary interest is to understand and quantify resistance during chemotherapy. For this, consistent with previous experimental [15, 16, 34, 38] and theoretical [7, 13, 21,22,37,39] studies of bet-hedging, we constructed a mathematical model of phenotypic switching to track the density of drug-sensitive (*A*(*t, a*)) and drug-tolerant (*B*(*t, a*)) cell phenotypes at time *t* and age *a*. We assume that cell phenotypes are fixed at birth [13], have phenotype-specific death rates *d_A_* and *d_B_*, and reproduce at rates 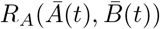 and 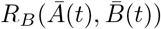, respectively.

Briefly, we consider multiple forms of 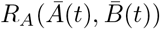 and 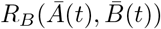 corresponding to different biological assumptions. When considering populations with (effectively) unlimited resources, such as those that are continually replated during *in vitro* experiments, we use a Malthusian growth model. We also consider the resource limited case, such as *in vitro* experiments that approach total confluence, and use a logistic growth model. Finally, we incorporate the effects of phenotypic cooperation, whereby a larger proportion of a certain phenotype can lead to increased phenotypic expansion through an Allee effect or frequency dependent fitness changes through a function 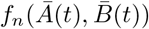 [2,40–43]. The function 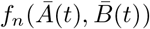 models the increase in relative fitness of drug tolerant cells as they become more common. Determining a precise formulation of 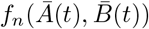 is difficult [23], and we give the functional forms of *R_A_* and *R_B_* in the Supplementary Information. Under these assumptions, *A*(*t, a*) and *B*(*t, a*) satisfy the age structured PDE,

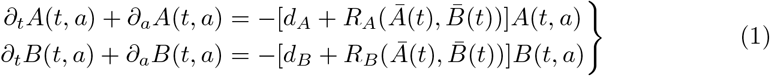

and the total number of cells of each phenotype at time *t* is calculated by

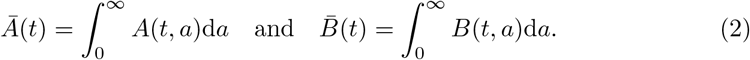

In this way, we studied the cellular ageing process over (linear) time (LHS Eq. (1)), with cellular loss at age *a* due to either death or reproduction (RHS Eq. (1)).

To model cellular reproduction, we assumed that the probability of changing phenotypes depended on the age of the parent cell (i.e., older cells are more likely to switch phenotypes during reproduction [44, 45]). The probability that a cell of age *a* and phenotype *i* will create a cell of phenotype *j* during reproduction is given by *β_ij_*(*a*), which leads to following boundary condition for Eq. (1).

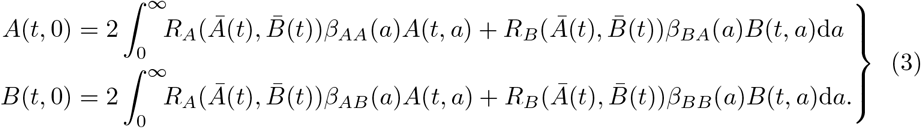

The probability of a cell with phenotype *A* and age *a* producing two daughter cells with the same phenotype is given by

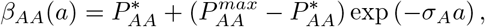

while the probability of a cell of phenotype *B* with age *a* producing two cells of phenotype *B* is

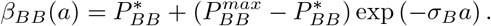

See Fig. S1 for representative forms and a discussion of *βAA* and *β_BB_*. Nascent cells were restricted to either phenotype *A* or *B*, i.e.

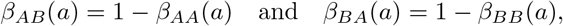

and we set

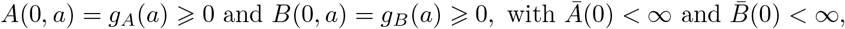

to be a biologically-relevant initial condition for the age distribution of cells (see the discussion in Supplementary Information for the technical conditions).

### Effects of phenotypic switching on population fitness

We first studied the role of phenotypic heterogeneity on population fitness in the presence of unlimited resources by considering two distinct measures of population fitness: the intrinsic growth rate of the population or Malthusian parameter λ*_P_*, and the expected number of offspring or basic reproduction number *R*_0_. In structured population models, these quantities are often related through the sign relationship: sign(λ*_P_*) = sign(*R*_0_ — 1) [46–48]. Precise mathematical formulations and results pertaining to these two metrics are described in the Supplementary Information.

A population comprised entirely of drug-sensitive cells has an intrinsic growth rate given by λ*_A_* = *r_A_* — *d_A_*, (and similarly, λ*_B_* = *r_B_* — *d_B_*, for a population of entirely drug-tolerant cells). The cost of resistance was assumed to reduce the intrinsic growth rate in the tolerant population, i.e., λ*_B_* < λ*_A_*. In a heterogeneous population of cells where cells cannot switch phenotypes, the Malthusian parameter is simply the maximum of growth rates of the constituent populations (λ*_P_* = max[λ*_A_, λ_B_*]). However, if cells exhibit phenotype plasticity, then the presence of a less-fit phenotype decreases the fitness of the combined population, and the intrinsic growth rate of the heterogeneous population falls between the growth rate of the constituent populations i.e., λ*_P_* ∈ (λ*_B_, λ_A_*) (Fig. 1). As we establish the previously mentioned sign relationship between *λ_P_* and *R*_0_ in the Supplemental Information, we can use either *λ_P_* < 0 or *R*_0_ < 1 as thresholds for population growth when designing a treatment schedule.

**Fig 1.**
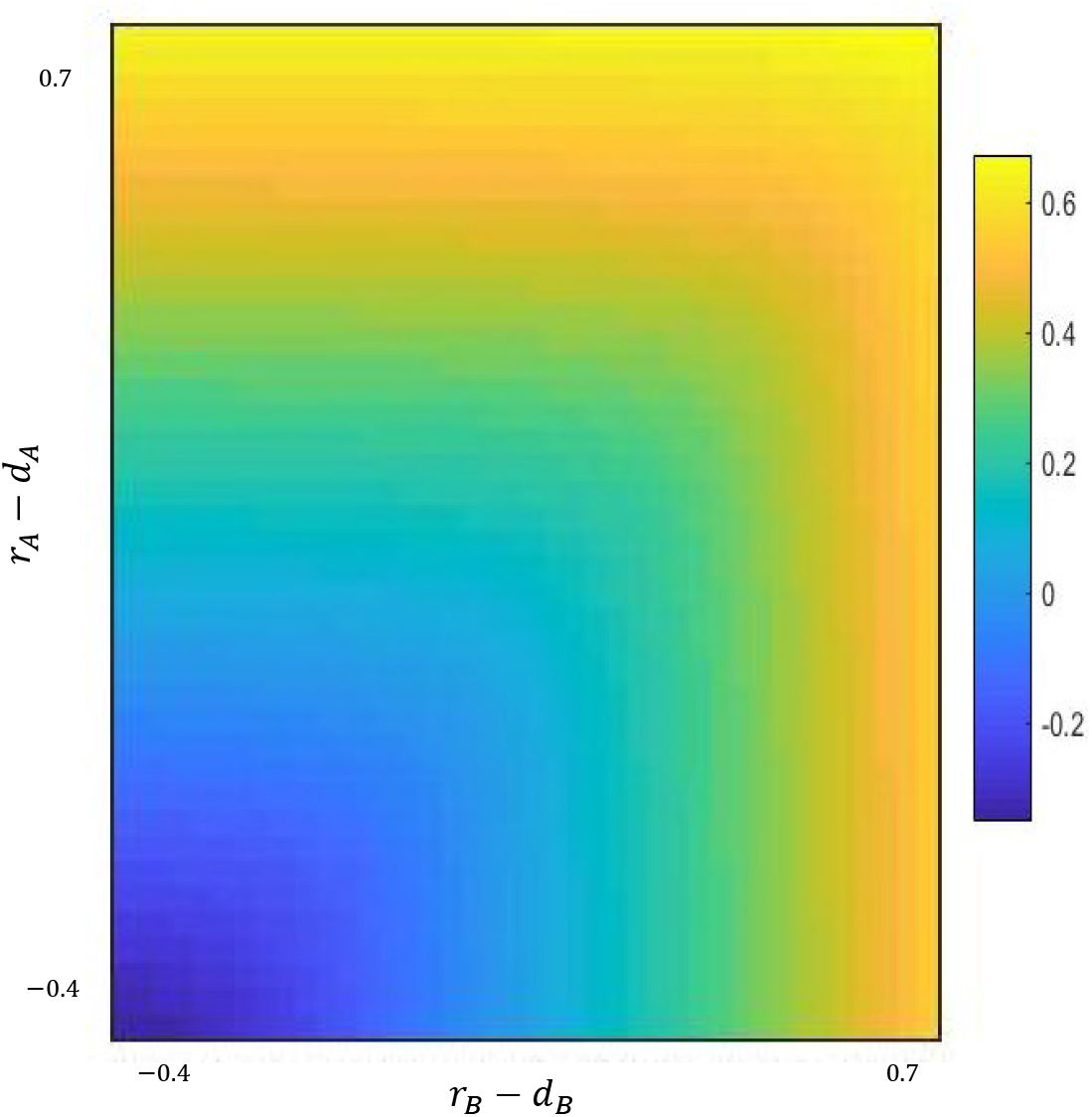
The population Malthusian parameter λ*_P_* as a function of intrinsic growth rates of sensitive and tolerant cells. The Malthusian parameter, λ*_P_*, is increasing along the main diagonal, and the figure is symmetric. Thus, increases in population fitness are driven by increases in the fitness of each phenotype without preference.

### Tumour composition evolves during population growth

In nutrient-rich environments, similar to serial replating in *in vitro* experiments, cooperation amongst drug-tolerant cells allows for tumour growth at a rate faster than purely Malthusian growth (Fig. S2). In the presence of unlimited resources, increasing the death rate *d_A_* of drug-sensitive cells while holding *r_A_* constant acts to decrease the fitness of the drug-sensitive cells (Fig. 2A and B), corresponding to both a decrease in the relative fitness difference between drug-sensitive and drug-tolerant cells and a decrease in the total population fitness. This is independent of the parameters that determine phenotypic inheritance (see Supplemental Information for details). We found that the proportion of sensitive type *A* cells was consistently higher during unlimited growth, in line with experimental results [20, 34]. Further, the strictly decreasing behaviour of the ratio 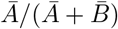 as *d_A_* increases indicates that the fitness difference between phenotypes plays a critical role in determining population composition during Malthusian growth.

**Fig 2.**
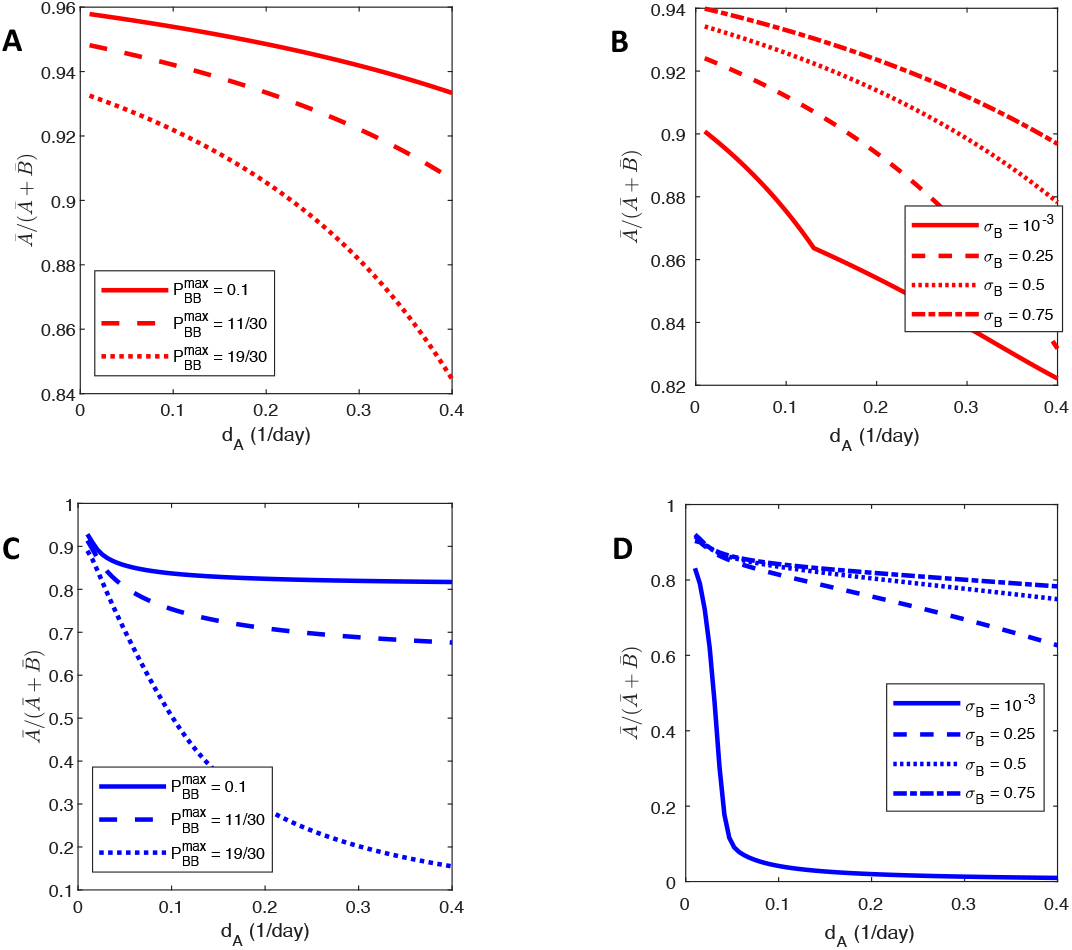
The ratio of drug-sensitive cells to the total population in Malthusian and resource limited growth for increasing values of the tolerant cell death rate. **A:** ratio *Ā*(*t*)/*N*(*t*) during Malthusian growth for 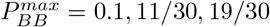. **B:** ratio *Ā*(*t*)/*N*(*t*) for *σ_B_* = [1 × 10^-3^, 0.25, 0.5, 0.75]. **C** and **D:** predictions as in **A** and **B** but in limited-resource settings. Note that the proportion of drug-tolerant cells in **A** and **B** compare favourably with the population composition of drug-tolerant cells reported by Sharma et al. [15].

In the limited resource situation, contrasting to the Malthusian case, the ratio 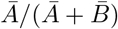 initially decreases before reaching a plateau and remaining relatively constant as d_A_ is increased. Accordingly, the relative fitness difference between phenotypes is less important that the probability of phenotypic switching in determining population composition (Fig. 2C and D). In fact, if the maximal probability of drug tolerant cells retaining their phenotype 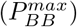 is sufficiently small, increasing *d_A_ increases* the proportion of type *A* cells (not shown). Conversely, if drug-tolerant cells are likely to produce drug-tolerant cells via a high probability of phenotypic inheritance, illustrated by the 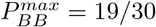 and 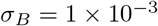 cases, the population evolves towards being predominantly drug-tolerant (phenotype B), despite the fact that λ*_A_* > λ*_B_*. Contrary to the unlimited resource case, where the relative fitness between phenotypes is the determining factor, the approximately constant proportion of drug-sensitive cells in the resource-limited setting suggests the importance of resource constraints in driving the establishment of a drug-tolerant population.

### Periodic Treatment Leads to Dominant Phenotype Switches

We next sought to quantify the permanence of treatment resistance. For this, we defined a phenotypic *switching strategy* as a pair 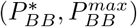 representing the minimal and maximal probability that a daughter cell retains the drug-tolerant phenotype of the parent cell. We considered two contrasting and illustrative switching strategies: 1) the *switching strategy* where resistant cells had a high probability of retaining their phenotype if they reproduced early in life (where this probability decreased to 0 as cells age, so 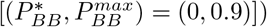, and 2) the *staying strategy*, where resistant cells were assumed to be unlikely to change phenotype regardless of reproductive age (i.e. 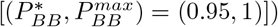. To measure the effectiveness of a given treatment strategy *S* for treatment from *t* = 0 to *t* = *T_end_*, we calculated the average total number of cells as a fraction of the carrying capacity,

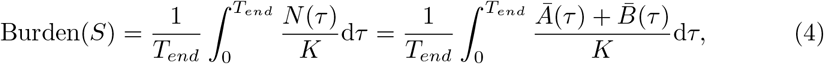

assuming here the physiologically-realistic finite resource case with phenotypic cooperation. Since we wish to avoid competitive release of the resistant sub-population, and subsequent tumour rebound, and are primarily interested in a sustainable reduction in tumour size, we consider the cumulative tumour burden over the entire treatment interval. In particular, (4) is related to the objective response rate, as schedules with a lower tumour burden as defined in (4) would presumably also have a higher objective response rate. Further, we used data from *in vitro* growth assays to parametrize our mathematical model. In these *in vitro* experiments, the population size can be easily measured and used as a proxy for treatment efficacy. However, if we were fitting the model to clinical data rather than *in vivo data*, it would be possible to use more clinically relevant measurements of treatment effect, such as time to disease progression or treatment failure due to resistance. In our framework, treatment resistance was defined as a significantly decreased therapeutic effect on the total population. The robustness of our results when considering different switching strategies is shown in the Supplementary Information.

We simulated a 21-day cyclic chemotherapy with the half-life of the anti-cancer agent set to *t*_1/2_ = 6 hrs, similar to cyclophosphamide, etoposide, and teniposide [49]. For both the *switch* and *stay* strategies, the tumour population eventually developed resistance as the drug-tolerant phenotype became dominant (Fig. 3). We observed that the proportion of drug-sensitive cells in the *switch* population remained above 20% of the total population – at least during the simulated treatment regimen – while the drug-sensitive cells in the *stay* population were effectively driven extinct during treatment. Thus different switching strategies act to either maintain or destroy a treatment-susceptible population. However, in the long term, the *switch* population eventually reverted back to a predominantly drug-sensitive population after treatment was discontinued. Clinically, this corresponds to transient resistance and an eventually re-sensitised population that has been observed in some cancers [8, 15, 16], suggesting that treatment holidays where therapy is re-applied after a break may be beneficial, as the *switch* population will eventually return to a mostly drug-sensitive state.

**Fig 3.**
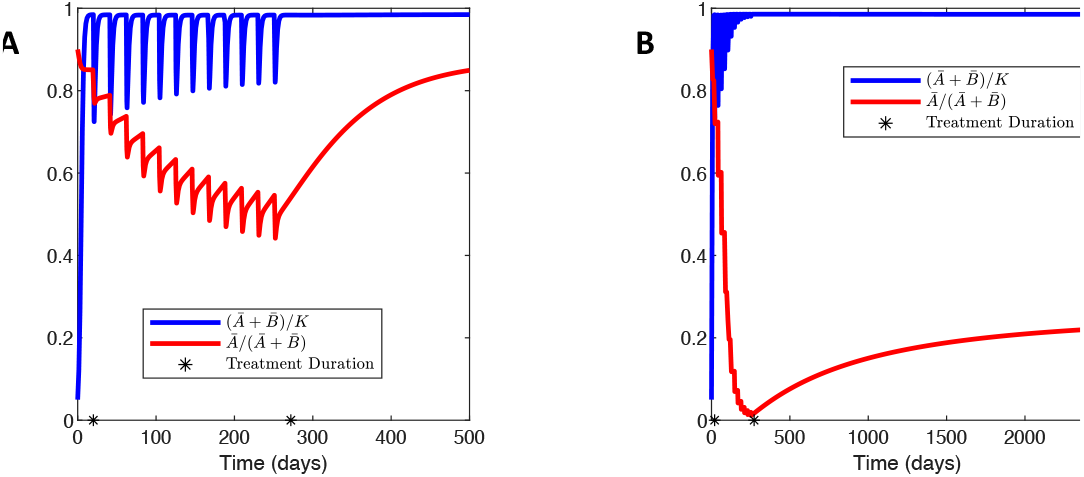
The effect of switching strategy on treatment efficacy. The effect of periodic treatment on a population using either a **A:** *switch* or **B:** *stay* strategy. In both cases, the same therapeutic strategy is applied between the black stars. In both **A** and **B**, the red curve shows the proportion of sensitive cells 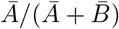, and the blue curve shows the dynamics of the total population *N*(*t*).

Conversely, the *stay* population evolved into a drug-tolerant phenotype dominated state and acquired essentially permanent resistance to therapy. In this case, reapplying the same therapy would be unsuccessful, even after a treatment holiday. These contrasting results demonstrate that different strategies of phenotypic switching can account for two drastically different types of therapeutic resistance by inducing either transient or permanent changes to the population, further underlining the difficulty of designing effective therapies to prevent phenotypic switching.

### Avoiding resistance to therapy in NSCLC

Lung cancer is the leading cause of cancer-related death in the United States, and non-small cell lung cancer (NSCLC) accounts for 20% of all cancer-related deaths [50]. Nearly two-thirds of NSCLC patients present with surgically unresectable disease and rely on systemic therapies for survival. Better characterisation of NSCLC at the molecular level has resulted in the introduction of a number of targeted therapies that are safer and more effective than standard chemotherapy. However, successful long-term treatment of NSCLC remains hampered by drug resistance [51]. We have recently shown that phenotypic interactions in co-cultured NSCLC spheroids and heterogeneity within patient samples drive a Cooperative Adaptation to Therapy (CAT) [7]. We sought to further quantify the evolution of phenotypic switching in NSCLC using our switching model to better understand treatment failure due to drug tolerance. In the subsequent analysis, we derived a number of analytical results and expressions that characterise our model informed treatment schedule. These quantities and their biological interpretation are summarized in Table 1 with the full analytical details given in the Supplemental Information.

**Table 1.** Summary of analytical expressions used to determine model informed therapy.

| Parameter | Value | Biological interpretation | Reference |
| --- | --- | --- | --- |
| $\lambda_A$ | $r_A - d_A$ | Intrinsic growth rate of drug-sensitive cells | Prop. S2 |
| $\lambda_B$ | $r_B - d_B$ | Intrinsic growth rate of drug-tolerant cells | Prop. S2 |
| $\lambda_A^*$ | $r_A - d_A^{max}$ | Treated growth rate of drug-sensitive cells | Definition |
| $\lambda_B(\theta)$ | $r_B f_n(\theta) - d_B$ | Growth rate of drug-tolerant cells<br>with Allee effect | Eq (7) |
| $\varepsilon$ | $\varepsilon \in (\frac{r_B}{r_A}, 1)$ | Permissible fitness from Allee effect | Chosen |
| $\vartheta_\varepsilon^*$ | $\left[ \frac{\varepsilon r_A - r_B}{r_A(1-\varepsilon)} \right]^{1/n}$ | Ratio $\bar{A}/\bar{B}$ to ensure $r_B < \varepsilon r_A$ with Allee effect | Eq (5) |
| $Dose^*$ | $\alpha_T \frac{\lambda_A}{-\lambda_A^*}$ | Minimum dose size to ensure $R_0^* < 1$ | Eq (6) |
| $\theta^*$ | $\left[ \frac{-\lambda_B}{\lambda_A} \right]^{1/n}$ | Threshold $\theta$ such that $\lambda_B(\theta) < 0$ | Eq (8) |

Beyond the intrinsic heterogeneity within tumours, external factors, including maximally tolerated dosing schedules, can lead to the establishment of a resistant phenotype and limit the effectiveness of therapy. This is clearly clinically disadvantageous. However, if there were no cooperation amongst drug-tolerant cells, then once the selection pressure of therapy was removed, the population would become re-sensitised to therapy. Thus, a possible therapeutic strategy is to limit the fitness gain of drug-tolerant cells due to cooperation. In our model formulation, limiting fitness gain due to cooperation is equivalent to limiting cooperation driven increases in reproduction rates, that is, *r_B_* < ε*r_A_*, where ε ∈ (*r_B_*/*r_A_*, 1). To accomplish this, the ratio of drug-tolerant to drug-sensitive cells, 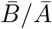, must not exceed the threshold ratio 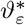 which is given by

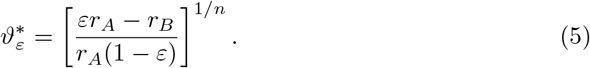

Using 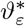, it is possible to schedule therapy to avoid competitive release (“fall then rebound”) as drug sensitive phenotypes switch to drug tolerant ones, and maintain a drug-sensitive population (Supplementary Information and [22,23]). Here, we detail a strategy for our NSCLC data that simultaneously ensures that the total tumour population decays and the population of drug-tolerant cells remains dependent on the drug-sensitive cells for survival. This strategy requires a delicate balance of maintaining chemotherapeutic concentrations at a large enough value to inhibit growth of the drug-sensitive cells while maintaining the frequency of drug-tolerant cells below a level that induces significant cooperation and the resulting competitive release. In the analytical work that follows, we assumed that *r_B_* < *d_B_*, an assumption which is satisfied by the results of the parameter fitting described in the Supplementary Information, and that there were ample resources available, though we include the carrying capacity in our simulations. Lastly, we assumed that the chemotherapeutic infiltrated the tumour uniformly and that therapy is administered over a fixed period *T*. We calculate the chemotherapeutic concentration during metronomic therapy in the Supplementary Information along with the precise analytical results underpinning our strategy.

From a classical population dynamics perspective, if the treated basic reproduction number 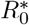 is less than 1, then the tumour population is expected to decay during treatment. In a approximately periodic environment, where chemotherapy is administered every *T* days, the threshold minimum dose size to ensure that 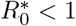 is

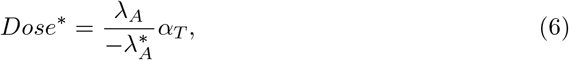

where 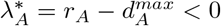 is the decay rate of the drug-sensitive population during treatment and *α_T_* is a constant depending on the period of administration *T*. We give a derivation of (6) and the explicit expression for *α_T_* in the Supplemental Information. To render this threshold clinically relevant, we rewrite Eq.(6) as

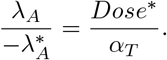

The left hand side above is comprised of patient specific parameters, namely the intrinsic growth rate of the drug susceptible population λ*_A_*, and the decay rate of the sensitive population during treatment, 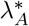. To estimate these quantities, consider two time series, *Ā_i_* and 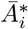, representing a drug-sensitive population grown in normal media or in the presence of a chemotherapeutic, respectively. Due to phenotypic switching, it is unlikely that these populations are comprised of solely drug susceptible cells, which complicates the estimation of λ*_A_* and 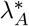 directly from experimental data. Nevertheless, to first approximation, the slope of log(*Ā_i_*) during the early exponential stage of growth offers an estimate for the intrinsic growth rate λ*_A_* = *r_A_* — *d_A_*. Assuming, for simplicity, that the chemotherapeutic agent only acts to increase the death rate of cells, then the the slope of 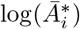 during exponential decay rate gives an estimate for 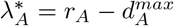. The right hand side of Eq (6) is comprised of parameters describing the properties of the drug as well as the size and frequency of drug administration that can be directly translated to the clinic.

To inhibit competitive release (i.e. the observed “fall and rebound” in the NSCLC spheroid data during therapy), we updated our approach to avoiding the establishment of tolerant phenotypes to ensure that λ*_B_* < 0 even when considering cooperation. Denoting the ratio of drug-tolerant to drug-sensitive cells by 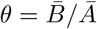, the expression for the fitness of the drug-tolerant population including the Allee effect is

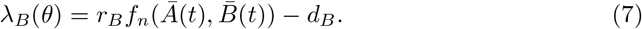

where 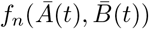 models cooperation mediated fitness increases and is given in Eq (S9). In particular, 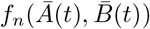 is a Hill type function with Hill coefficient *n* ⩾ 1. In this context, n can be understood as representing the necessary amount of cooperation between drug-tolerant cells to induce a fitness increase. Then, enforcing the condition that λ*_B_* < 0 yielded the threshold ratio

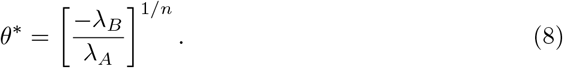

As λ*_B_* < 0, the right hand side of Eq (8) is the ratio of the decay rate of drug-tolerant cells to the growth rate of drug-sensitive cells. If the population of drug-tolerant cells decays at a faster rate than the population of drug-sensitive cells grows (|λ*_B_* | > λ*_A_*), then drug-tolerant cells must outnumber drug-sensitive cells before cooperation will allow for expansion of the drug-tolerant population. In this case, cooperation acts to attenuate the factor by which drug-tolerant cells must outnumber drug-sensitive cells before the drug-tolerant population is self-sustaining, since

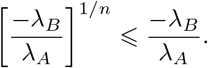

Conversely, if |λ*_B_* | < λ*_A_*, then increasing levels of cooperation necessitates a larger proportion of drug-tolerant cells to permit self-renewal of the treatment resistant population. Once again, the ratio of untreated intrinsic growth rates can be directly estimated from *in vitro* data. In summary, ensuring that *θ* < *θ*\* is sufficient to avoid the establishment of a resistant population.

### Model-informed treatment drives tumour extinction

Lastly, we combined the strategies ensuring tumour decay or avoiding the establishment of resistance described above to drive long-term treatment effectiveness to docetaxel (see Supplementary Information for similar results for the chemotherapeutics afatanib and bortezomib and details of the parametrization of the pharmacokinetic models for each therapeutic).

In most treatment schedules, docetaxel is administered either weekly or once-every-three-weeks [52], however, it is not obvious that either of these cycle lengths represent optimal treatment periods. Rather, as suggested by Bacevic et al. [23] and others, it may be ideal to dose more frequently and with less intensity to maintain drug pressure on the population. Therefore, we did not *a priori* fix the period of administration *T* to model-informed therapy. Rather, for *T* = 1, 2, 3…, 7 days, we determine the model-informed dose size as

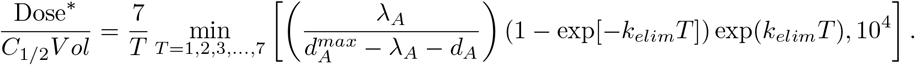

Increasing the density of therapy increases the burden of therapy and may be overwhelmingly toxic. Accordingly, we imposed that the cumulative chemotherapeutic dose under model informed therapy does not surpass what would be administered in the fixed periodic schedule. For each period *T*, we used the minimal dose size that satisfies (6), as our calculation of 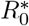 only identifies a sufficient condition to drive tumour population decay, and any larger dose size potentially allows competitive release of the drug-tolerant population.

We tested the model-informed therapy for each value of *T* = 1, 2, 3, …, 7 and chose the largest therapy period *T* that avoided the establishment of a drug-tolerant population and led to sustained population decay. While the decision to administer therapy on each treatment day *nT* is dependent on the ratio *θ*(*t*) < *θ*\*, and the tumour micro-environment may not be precisely periodic, our results indicate that combining the two model informed constraints successfully drives tumour extinction.

We compared our model-informed therapy to periodic dosing administered every 7 days and found that informed therapy performed comparatively to periodic dosing during the initial stage of therapy (Fig. 4). However, the benefit of our model-informed therapy becomes apparent when inspecting the behaviour of the treated tumour over longer periods: the fixed dosing schedule allowed for the establishment of a drug-tolerant phenotype and the eventual loss of effectiveness of therapy, while the model-informed therapy maintained a drug-sensitive population and led to sustained tumour decay.

**Fig 4.**
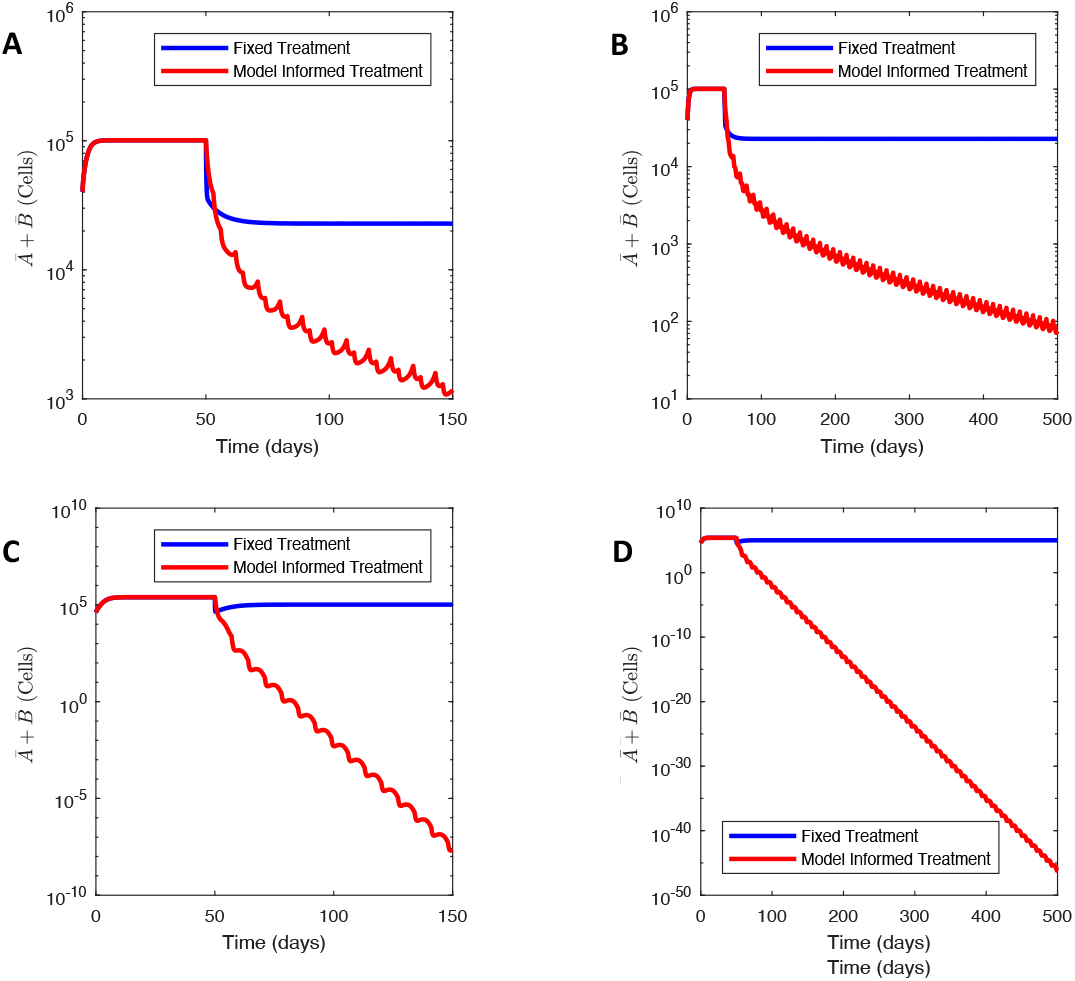
Comparing model-informed and periodic dosing treatment regimens. Panels **A** and **B** show the effect of model-informed therapeutic strategy on the WT population over 100 and 450 days, respectively with *T* = 3. Panels **C** and **D** show the effect of model-informed therapeutic strategy on the M1 population over 100 and 450 days with *T* = 7, respectively. The parameters used in this simulation are given in **S3** TableS1.

As before, we computed the effectiveness of therapy using Eq. (4). The ratio of tumour burden in the model-informed therapy to periodic therapy was 0.2059 and 0.3708 for Wild Type (WT) and Mutant 1 (M1) cells studied (see *Materials and methods*), respectively. This result clearly demonstrates that model-informed therapy significantly outperforms periodic therapy, and is consistent across all populations and therapeutics, demonstrating the robust efficacy of this adaptive approach to maintaining tolerant phenotypes.

## Discussion

Despite the introduction of novel targeted therapies and increased characterisations of individual patient’s genetic landscapes, drug resistance continues to drive treatment failure. This suggests that identifying and understanding non-genetic factors contributing to drug therapy tolerance is crucial to providing better care. In this work we proposed a simple quantitative model of stochastic phenotype switching in the context of cancer. Our model is comprised of two non-local age structured PDEs that incorporate phenotypic switching through non-local boundary terms. Specifically, phenotypic switching is described as a random process where the probability of inheriting the parents’ phenotype is a decreasing function of cellular age at reproduction. This mapping from age to switching probability generalises the role of molecular switching mechanisms and the inheritence of signalling factors in phenotype determination. In this sense, we have studied the role of phenotypic ‘memory’ governed by the inheritance of intracellular factors, similar to the biological phenomenon observed by Yang et al. [31] where inheritance of signalling factors such as p53 and mitogen can predispose daughter cells towards quiescence or proliferation. Recent experimental work has implicated these signalling factors and the resulting non-genetic memory in response to anti-cancer treatment. These results have specifically identified the role of treatment induced stress on mother cells as a determining factor in daughter cell’s adoption of a “persister” like phenotype [53–55]. In previous theoretical work we demonstrated that the precise mechanisms governing phenotype switching determine the rate of extinction under cytotoxic therapy [13]. Thus, molecular switching mechanisms may be subject to evolution by natural selection. The model presented here represents a more general framework to further explore this phenomenon, and to extend it through the introduction of phenotypic memory, by specifying switching dynamics in a functional, rather than network-defined, form.

Given the assumption of unlimited resources, we derived expressions for the Malthusian parameter and basic reproduction number, and established the classic sign relationship between these two measures of population fitness. From this model, we derived an equivalent ODE model describing the dynamics of drug-tolerant and sensitive populations to study the impact of resource availability and intra-phenotype cooperation on population growth. This allowed us to show that competition for limited resources facilitates the establishment of a less fit phenotype. Incorporating a phenomenological model of cytotoxic therapy, we showed that the phenotype switching strategy of the drug-tolerant population (to either preferentially inherit or relinquish the parent cells phenotype) determined the type of treatment resistance observed. In particular, our mathematical model can reproduce both transient drug resistance or epigenetic permanent resistance by only changing the switching strategy of the drug-tolerant population. Leveraging this, we proposed a treatment schedule that exploits the population composition to avoid the establishment of treatment resistance.

Importantly, we then applied our model to understand the development of treatment resistance within *ex vivo* NSCLC tumour spheroids to understand the impact of phenotypic switching on response to treatment. When exposed to chemotherapeutics, genetically identical NSCLC populations were found to exhibit a “fall then rebound” behaviour indicative of phenotypic resistance. We derived the basic reproductive number in the context of periodic treatment and determined a therapy schedule that avoided the establishment of resistance and exhibited sustained tumour decay. The NSCLC data and our results underline that phenotypic switching may be occurring in a genetically identical population of NSCLC cells and may be driving treatment resistance. It is thus important to quantify it’s presence and impact of on treatment scheduling. In this work, we presented a mathematical model to understand phenotypic heterogeneity and derived a model informed strategy to mitigate –and potentially avoid–phenoytpically driven treatment resistance.

Our phenomenologically-based model is simple. Consequently, our results must be evaluated in light of the many assumptions and limitations of our model, and remain to be further validated in experimental systems. We also made an important assumption that cancer cells are either entirely drug-tolerant and drug-sensitive. While this assumption of a discrete phenotype landscape simplifies the mathematical modelling, it is not biologically realistic. Moreover, we do not consider the role of spatial and metabolic heterogeneity [56–58], drug infiltration [59,60], nor the role of other cells in the tumour micro-environment [61].

These limitations notwithstanding, our work identifies the role of stochastic switching in therapeutic resistance, explicitly incorporates non-genetic inheritance, or phenotypic memory, in a physiologically structured mathematical model, and highlights the role of mathematical modelling in understanding and developing evolutionary-inspired therapeutic strategies.

## Materials and methods

### Non-small cell lung cancer data

We integrated *in vitro* growth assay data taken from a NSCLC cell line with induced mutations in Dicer1 [7]. Briefly, Craig et al. [7] cloned a cell line with an oncogenic Kras, homozygous p53 and heterozygous Dicer1 loss of function mutations that induces tumours when injected into mice. Growth of non-small cell lung cancer (NSCLC) tumour spheroids was quantified as previously described [7, 62]. The parental (WT) cell line was derived from KRas-G12D, p53—/—, Dicer1f/— genotype lung tumours and mutants (M1 and M2) were obtained through transfection to Dicer1+/+ and Dicer1-/- using CRISPR-Cas9 [62]. Cells were then transfected with lentivirus particles expressing fluorescent proteins for quantification using flow cytometry [7]. Cells were plated as tumour spheroids on NanoCulture plates every other day for 7 days in the absence or presence of a variety of anti-cancer agents [7]. WT, M1, and M2 were all grown separately as monocultures and in co-culture as parental and mutant lines in proportions of 10:90, 50:50, and 90:10. Growth without and with drug was assessed via flow cytometry on days 1, 3, 5, and 7 [7].

In the untreated experiments, Craig et al. [7] cultured a genotypically homogeneous population of NSCLC cells in a constant environment for 7 days. In the treated experiments, after 72 hours of growth in untreated medium, the authors bathed the population of cells in a constant and lethal concentration of one of three chemotherapeutics (docetaxel, afatinib, or bortezomib) and counted the number of surviving cells. As the anti-cancer drug concentration is constant, we assume that the observed “fall and rebound” behaviour is not driven by the proliferation of a drug-sensitive population, but rather due to the expansion of a drug-tolerant population, similar to the phenotypic resistance observed in numerous studies [8, 15, 16]. While the drug-tolerant population may have arisen due to genetic mutations, the short treatment time of 96 hours suggests the expansion of a previously established drug-tolerant phenotype. We report results for the WT and Mutation 1 (M1) lineages treated with docetaxel in the main text with similar results for WT, M1 cells treated with afatinib and bortezomib, as well as a separate population Mutation 2 (M2) cells shown in the Supplemental Information.

### Numerical simulation of phenotypic switching model

Equation (1) is a system of coupled non-local PDEs for the cell densities *A*(*t, a*) and *B*(*t, a*). Rather than implementing these PDEs numerically, we note that we are primarily interested in the number of drug-sensitive and drug-tolerant cells, given by

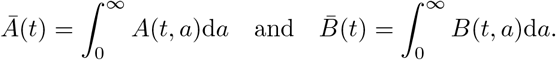

For *L_1_* initial data, the theory of transport equations ensures that these integrals are finite [63]. Therefore, rather than solving the system of coupled PDEs and integrating over age to compute *Ā*(*t*) and 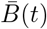, we derive an equivalent finite dimensional system of ordinary differential equations for the populations *Ā*(*t*) and 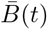. The derivation uses Leibniz’s integral rule and integration by parts and is detailed in the Supplemental Information. Incorporating phenotypic switching through the boundary conditions necessitates two extra ODEs for the proportion of drug-sensitive or drug-tolerant cells retaining their phenotype. The resulting ODE model is

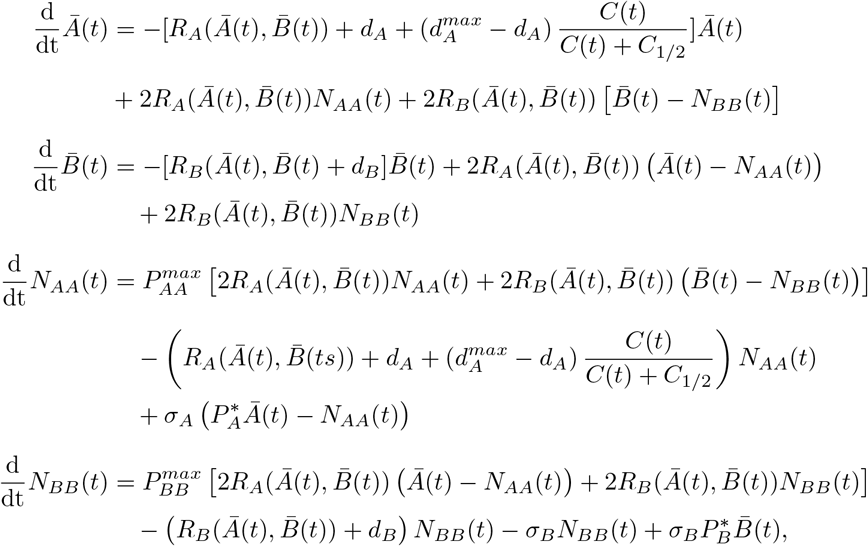

with initial conditions corresponding to populations in exponential growth.

### Generic model of chemotherapy

We denote the concentration of a chemotherapeutic at time *t* as *C*(*t*) and assume that therapy is given intravenously with administrations at times 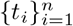. The time dynamics of *C*(*t*) are given by

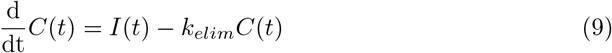

where *I*(*t*) models the I.V administration of the cytotoxic drug during an injection time of *T_admin_*, and is given by

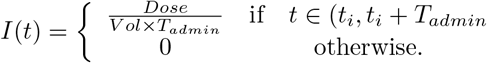

and where *V ol* is the volume of absorption of the drug and *Dose* is the size of each administration. The half life of the drug in question, *t_1/2_*, defines the elimination constant through *k_elim_* = log(2)/*t*_1/2_.

We assume that chemotherapy increases the death rate of drug-sensitive cells through

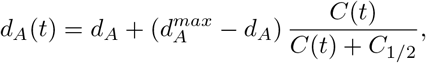

where the half effect concentration is given by *C*_1/2_. We note that, in our simple model, it is the ratio of the drug concentration *C*(*t*) and the half effect *C*_1/2_ that completely determine the pharmacodynamics of the therapy in question. While using this simple pharmacodynamic model limits the direct applicability of our work, it allows for the identification of the crucial aspects in determining the effect of therapy.

### Model parametrization to NSCLC data

To fit the mathematical model to the NSCLC *in vitro* data, we fix 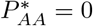 and 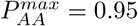, set σ*_A_* = σ*_B_* = 1 × 10^-2^ hours^-1^, and enforce *d_A_* = *d_B_* and account for the fitness cost of resistance by enforcing *r_B_* ⩿ *r_A_*. The parameters remaining to be fit control either population growth (*r_A_,r_B_*, and *d_A_*), or the probability of retaining the drug-tolerant phenotype (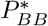 and 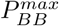).

We show in the Supplemental Information that, for given untreated data, these parameters may not be identifiable. In particular, for a given pair 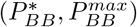, it is possible to fit the parameters *r_A_, r_B_* and *d_A_* can be chosen to fit experimental data equally well in the absence of treatment. However, the role of the parameters 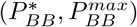 becomes evident once therapy is administered and the previously indistinguishable curves become distinct. Therefore, we simultaneously fit the untreated and docetaxel data from [7].

During the treated experiments, the cells are continuously bathed in lethal concentrations of each chemotherapeutic, so we model the death rate of the drug-sensitive cells during fitting as

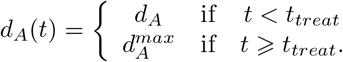

For treated and untreated time series data 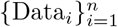, we fit the parameters 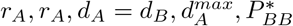 and 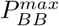 by minimizing

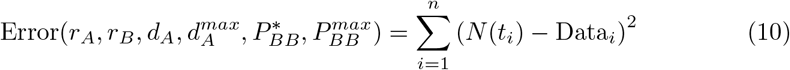

where 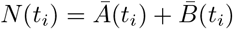 is the total number of cancer cells predicted by the mathematical model. We used the Matlab [64] algorithm *fmincon* to minimize (10) with 15 initial starting points in parameter space. The results of our fitting to the untreated and docetaxel data are shown in Fig. S5.

Having fit the parameters 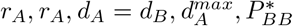 and 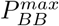 to the untreated and docetaxel treated population data, we fix the tumour growth parameters *r_A_,r_A_,d_A_* = *d_B_*, and only fit 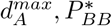 and 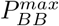 for the data from experiments with afatinib and bortezomib. We do not refit the growth rate *r_B_*, both to avoid overfitting, and as cancer cells with a drug-tolerant phenotype have exhibited cross-resistance to other chemotherapeutics [16]. We list the tumour growth and switching parameters in Tables S1 and S2.

## Supporting information

**S1 Text Supplementary information file**.

**S1 Fig. Phenotypic switching probability and relative fitness gain**. Panel **(A)** shows a representative form of *β_ii_* **(A)**. Panel **(B)** shows the frequency dependent fitness increase function *f_n_*(*θ*) for *n* = 1, 2, 10.

**S2 Fig. A comparison of growth rates for different growth functions** *f_n_, n* = 1, 2, 3,10, **against Malthusian growth**. The “no fitness” curves corresponds to no frequency dependent fitness increase and *f_n_* = 1. Panel **(A)** shows population evolution from an initial population comprised of 100 drug-sensitive cells and one drug-tolerant cell. Conversely, Panel **(B)** shows population evolution from an initial population comprised of one phenotype *B* cell, or 1 drug-sensitive cell and 100 drug-tolerant cells.

**S3 Fig. Fitting of the mathematical model to the Dingli 2009 data for a variety of fitting strategies**. Panel **(A)** shows the fitting of the mathematical model (1) to the Dingli 2009 data for 8 different switching strategies. Panel **(B)** shows the same 8 strategies after 2 applications of therapy.

**S4 Fig. The effect of adjustable therapy on a partially resistant population** The drug-tolerant population uses either a *switch* (panel **(A)**) or *stay* (panel **(B)**) strategy. In both cases, treatment is applied between the black stars, while the red curve shows the proportion 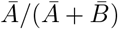, and the blue curve shows the dynamics of *N*(*t*).

**S5 Fig. Fitting results of Equation** (1) **to the WT and M1 data treated with docetaxel** The data for WT and M1 are given in Panels **(A)** and **(B)**, respectively. In all cases, the untreated data is given by the black stars while the untreated simulation is in solid blue. The docetaxel treated data is given by the hollow circles and the treated simulation is in dashed blue.

**S5 Fig. Fitting results of Equation** (1) **to the WT, M1 and M2 data from treated with afatinib and bortezomib**. Panels **(A)**, **(B)** and **(C)** show the WT, M1 and M2 data treated with afatinib, respectively. Panels **(D)**, **(E)** and **(F)** show the WT, M1 and M2 data treated with bortezomib, respectively.

**S1 Table The switching parameters for WT, M1, and M2 cell lines**.

**S2 Table The tumour growth parameters for the WT, M1 and M2 type cells**.

**S3 Table The effectiveness of model informed therapy when compared to periodic dosing over 150 days**.

**S6 Fig. Comparison of periodic therapy in blue and model informed therapy in red for afatinib and bortezomib**. Panels **(A)** and **(B)** are the WT and M1 cells treated with afatinib, respectively. Panels **(C)** and **(D)** are the WT and M1 cells treated with bortemozib, respectively.

## Acknowledgements

Portions of this work were completed while TC, DN, and ARAA participated in the thematic semester in Mathematical Biology at the Institut Mittag-Leffler. TC is grateful for many useful conversations with Tony Humphries.

## Supplementary Information: The role of memory in non-genetic inheritance and its impact on cancer treatment resistance

### General model of phenotype switching

Here, we detail the mathematical model used in the main text. As mentioned, we consider two distinct cellular phenotypes *A* and *B*, where phenotype *A* represents the drug sensitive sub-population and phenotype *B* represents the drug tolerant sub-population. We denote the age density of cells with phenotype *A* at time *t* as *A*(*t, a*), while *B*(*t, a*) represents the density of cells of age a with phenotype *B* at time *t*. The object of clinical interest at time *t* is unlikely to be the density of cells with a given age, but rather total number of cells of each phenotype, given by

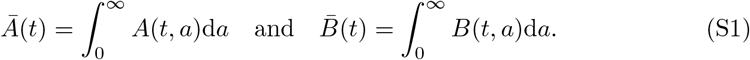

Under the assumptions in the main text, A(t, a) and B(t, a) satisfy the non-local PDE

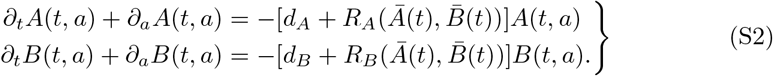

Cellular reproduction is incorporated through non-local boundary conditions. Specifically, reproduction of cells produces 2 daughter cells with age a = 0 that may not inherit the parent’s phenotype. As mentioned, we assume that the probability of changing phenotypes depends on the age of the parent cell: i.e, older cells are more likely to switch phenotypes during reproduction [44,45]. We use *β_ij_*(*a*) to denote the probability that a cell with age a and phenotype *i* will create a cell of phenotype *j* during reproduction. Hence, the boundary condition corresponding to (S2) is

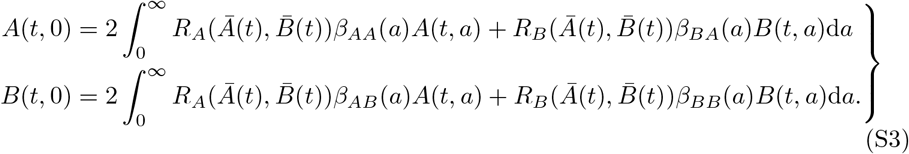

We model the probability that a cell of phenotype *A* gives birth to two cells of phenotype *A* as

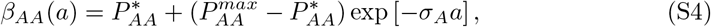

while the probability of a cell of phenotype B producing two cells of phenotype *B* is

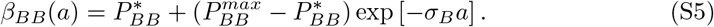

As expected, we note that both *β_AA_*(*a*) and *β_BB_*(*a*) are non-negative decreasing functions of age. Moreover, as nascent cells are assumed to be restrained to either phenotype *A* or *B*, we necessarily have

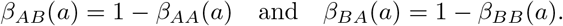

We illustrate a representative form of *β_ii_*(*a*) in Figure S1 (a). Using the bistable switch example, the σ*i* parameters model the decay rate of the molecules that bias the switch. With σ*i* = 1 × 10^-2^ days^-1^ (as is the case in our generic parametrization), a cell that replicates after 1 day will a roughly 1% smaller probability of phenotypic inheritance than if the cell had replicated immediately upon birth.

To complete the initial value problem defined by (S2), we prescribe initial conditions describing the age distribution of cells at time *t* = 0

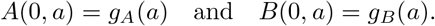

It is natural to enforce that the initial age distributions *g_A_* (*a*) and *g_B_* (*a*) are non-negative functions. We note that cells with age *a* > 0 at time *t* = 0 must have been born at some time *s* < 0. However, *A*(*t, 0*) and *B*(*t, 0*) are only defined for *t* > 0. Nevertheless, it is possible to use the initial data of (S2) to define *A*(—*s*, 0) and *B*(—*s*, 0) for *s* ∈ (0, ∞) by *A*(—*s*, 0) = *g_A_*(*s*) exp([*r*_B_ + *d*_B_]*s*) and *B*(—*s*, 0) = *g_B_* (*s*) exp([*r_B_* + *d_B_*]*s*) [65], and we use these definitions when considering the equivalent renewal equation (S25).

Further, we are only interested in finite populations, so we require that *g_A_,g_B_* ∈ *L_i_*(0, ∞). The requirement that *g_A_* and *g_B_* are integrable ensures that (S2) has a unique solution in the sense of distributions, and that this solution belongs to *L_1_* (0, ∞) [63]. It follows that the quantities *Ā*(*t*) and 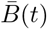 are well-defined for *t* > 0. Rather than introducing a maximal age and subsequent mathematical complications in our simple model, we note that, along the characteristics of (S2) given by a = *t* — *t*_0_,

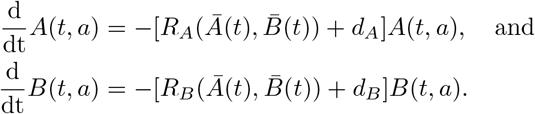

both *A*(*t, a*) and *B*(*t, a*) decay exponentially in age. Nevertheless, given the biological interpretation of a in Eq. (S2), it may be reasonable to enforce a maximal cellular age *a_max_*. However, translating this requirement to solutions of Eq. (S2) is not trivial.

Age structured PDE models similar to (S2) have been used extensively to model the progression of cells through a reproductive process [63, 65–68], and there is extensive mathematical theory regarding the use of these age structured models in mathematical biology (see [69] for a review). As mentioned in the Main Text, other authors have considered PDEs structured in phenotype with non-local or diffusion terms. However, to our knowledge, the incorporation of the phenotypic switching in a McKendrick type equation is new.

#### Growth dynamics

To complete the mathematical model (S2), we now specify the form of *R_A_* and *R_B_*. In what follows, we explore multiple forms of these functions corresponding to different biological assumptions. We begin with the simplest case: unconstrained exponential growth which is appropriate in populations with (effectively) unlimited resources such as those that are continually replated during *in vitro* experiments. We then remove this assumption of unlimited resources and consider constrained growth, such as *in vitro* experiments that approach total confluence. Finally, we incorporate the effects of phenotypic cooperation, whereby a larger proportion of a certain phenotype can lead to increase phenotypic expansion through an Allee effect or frequency dependent fitness changes [2, 40–43]. By considering the effects of different growth functions, we will explore the impact of growth stage on establishment of a drug tolerant population.

The simplest case corresponds to unconstrained growth, where there are unlimited resources in the environment. This surplus of resources allows for exponential, or Malthusian growth. In this case, we use a constant and phenotype dependent growth rate

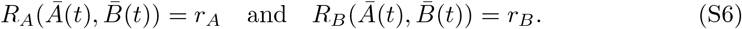

During *in vitro* experiments, this unconstrained growth corresponds to the early growth phase of cells in culture or following the replating of an established cell culture into a nutirent rich environment.

In an environment with limited resources, the early exponential growth of a population gives way to tempered growth as competition for resources begins. This restrained growth is typically modelled as logistic type growth. Thus, in the limited resource case, we model the growth rates as

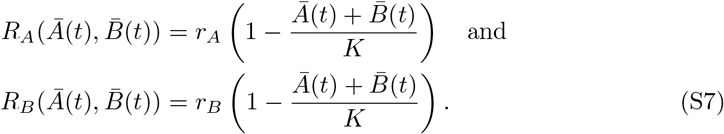

We note that, as the population reaches the carry capacity of the environment, the reproduction rates *R_A_* and *R_B_* both converge to 0.

Finally, we consider the influence of frequency dependent fitness increases in the *B* phenotype. This corresponds to cells of phenotype *B* gaining fitness as they become more populous and cooperate. This frequency dependent fitness increase, or Allee effect, has been observed in cancer [5, 6, 23,70–72]. To allow for cooperation amongst drug tolerant cells, we model reproduction as

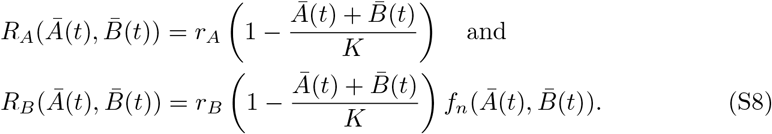

The function 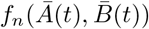 models the increase in relative fitness of drug tolerant cells and determining a precise formulation of 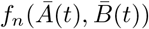 is difficult [23]. However, from biological considerations, as the proportion of drug tolerant cells increases, the relative fitness these cells should increase. Therefore, 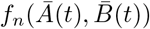 should be monotonically decreasing in *Ā* and increasing in 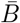. In what follows, we will use the following frequency dependent growth function

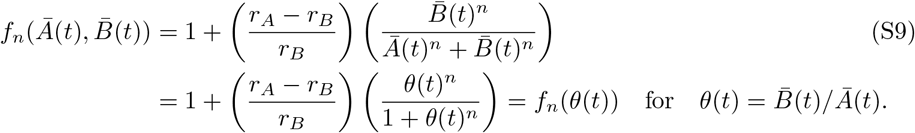

The function 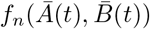 is a Hill-type function that ensures that 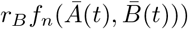 smoothly interpolates between *r_B_* and *r_A_* as the proportion of phenotype *B* cells in the total population increases between 0 and 1. The parameter *n* controls the steepness the sigmoidal curve. As *n* increases, the smooth sigmoid function approaches a step function at *θ*(*t*) = 1. Throughout the rest of this work, we consider *n* = 1, 2, 10 to illustrate the impact of different frequency dependent fitness functions on population dynamics. In Figure S1 (b), we show 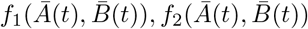 and 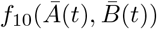 as a function of *θ*(*t*). We conclude by noting that the growth rates given by Equations (S6), (S7) and (S8) are non-negative functions for non-negative input *Ā*(*t*) and 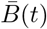.

#### Generic model of chemotherapy

Treatment necessarily imposes selection pressure against susceptible cells [5, 13, 14, 22, 23]. This selection pressure can drastically change population level dynamics and lead to the development and competitive release of resistant populations. As mentioned, this resistance can be driven by phenotypic switching [8, 15, 16]. Therefore, we include the effects of cytotoxic treatment in the mathematical model (S2). We emphasize that we are only attempting to model the qualitative effects of cytotoxic treatment and recall that, when incorporating therapeutic effects, we have assumed that

**Fig S1.**
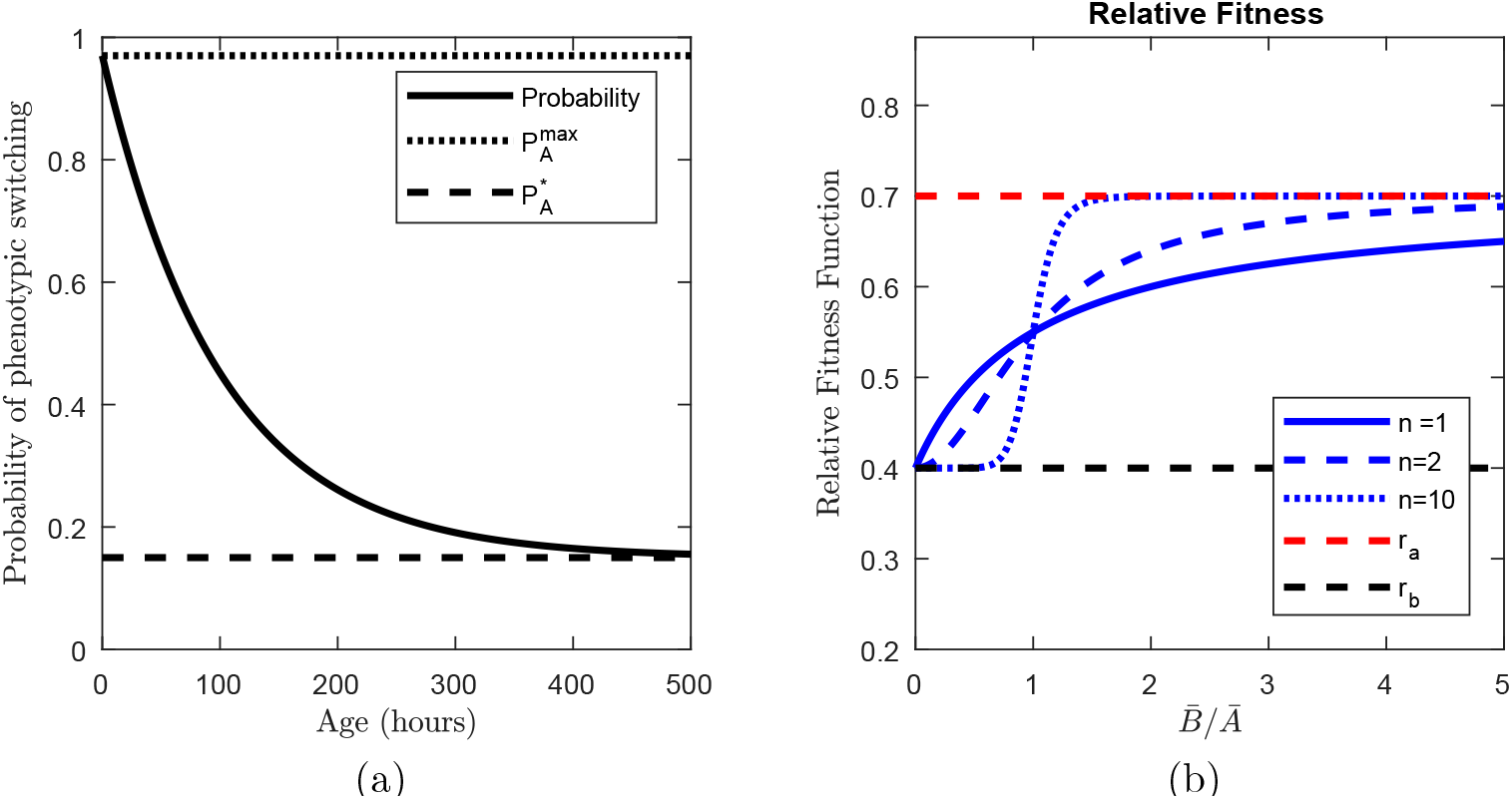
Phenotypic switching probability and relative fitness gain. Figure (a) shows a representative form of *β_ii_*(*a*). Figure (b) shows the frequency dependent fitness increase function *f_n_*(*θ*) for *n* = 1, 2, 10.

cells of phenotype *A* are drug sensitive, while cells of phenotype *B* are drug tolerant and thus resistant to treatment.

We denote the concentration of a chemotherapeutic at time *t* as *C*(*t*) and assume that therapy is given intravenously. The time dynamics of *C*(*t*) are given by

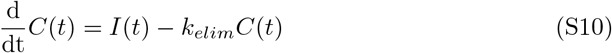

where *I* (*t*) is given by

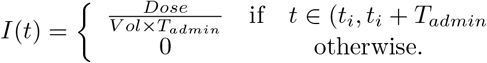

and models the I.V administration of the cytotoxic drug during an injection time of *T_admin_*, where *Vol* is the volume of distribution of the drug and *Dose* is the size of one administration. The half life of the drug in question, *t_1/2_*, defines the elimination constant through *k_elim_* = log(2)/*t*_1/2_.

We assume that chemotherapy increases the death rate of drug sensitive cells through

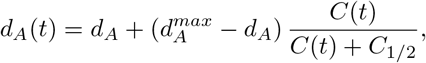

where the half effect concentration is given by *C_1/2_*. We note that, in our simple model, it is the ratio of the drug concentration and the half effect *C*_1/2_ that completely determine the pharmacodynamics of the therapy in question. While using this simple pharmacodynamic model limits the direct applicability of our work, it allows for the identification of the crucial aspects in determining the effect of therapy.

#### Ordinary differential equations for *Ā*(*t*) and 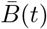

The partial differential equation (S2) is difficult to solve numerically. Moreover, we are primarily interested in the clinically relevant and biologically measurable *Ā*(*t*) and 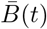. We note that the quantities of interest, *Ā*(*t*) and 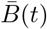, are independent of age. Therefore, it is reasonable to expect that their dynamics should be determined by a system of two ODEs. As we will show, the age dependence in the non-local boundary condition (S3) will necessitate the inclusion of two extra ODEs. As the analysis that follows is identical for 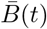, we only show the derivation of the ODE for *Ā*(*t*).

To derive the equivalent system of ODEs, we note that the formal solution of (S2) for A(t, a) during treatment is

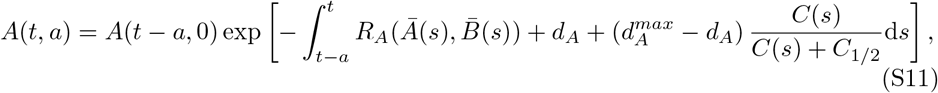

with a similar expression for *B*(*t, a*). For notational convenience, we denote the total population of cells at time *t*, 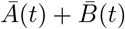 as *N*(*t*).

We begin by using Leibniz’s rule to differentiate (S1) and, after adding 0 = *∂_a_A*(*t, a*) — *∂_a_A*(*t, a*), find

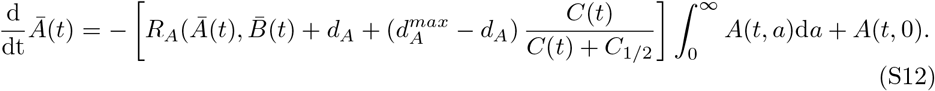

The boundary conditions of (S2) give

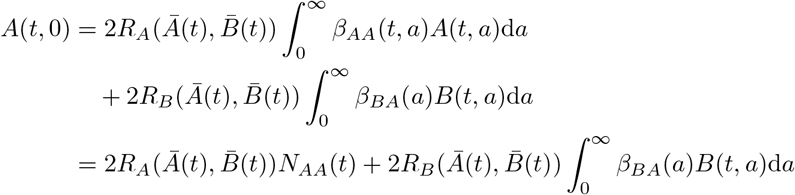

where

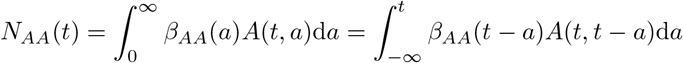

with a similar expression for *N_BB_*(*t*). We note that, as *A*(*t, a*) ∈ *L_1_*(ℝ_+_) with the normal Lebesgue measure, and *β_AA_* ⩽ 1, it follows that *N_AA_* is finite. Now, we recall that reproducing cells of phenotype *i* either create two cells of phenotype *i* or *j*. Therefore, so

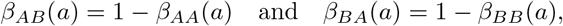

so

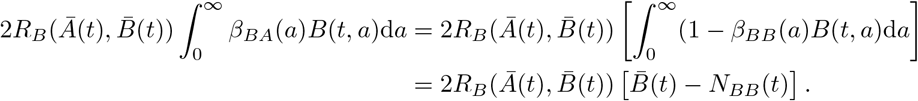

Thus, *Ā*(*t*) and 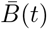 satisfy a system of differential equations that require the evaluation of the integral terms *N_AA_*(*t*) and *N_BB_* (*t*), which is, once again, numerically challenging. Therefore, to implement (S12) numerically, we write *N_AA_*(*t*) and *N_BB_* (*t*) as the solutions of the differential equations

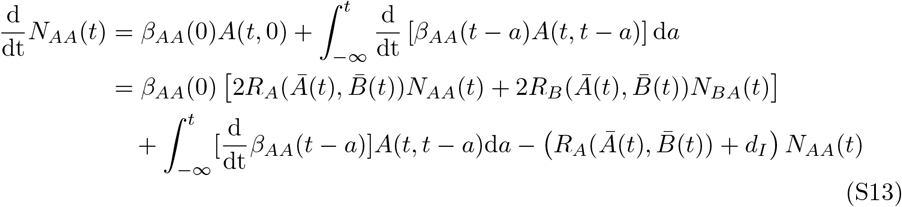

an

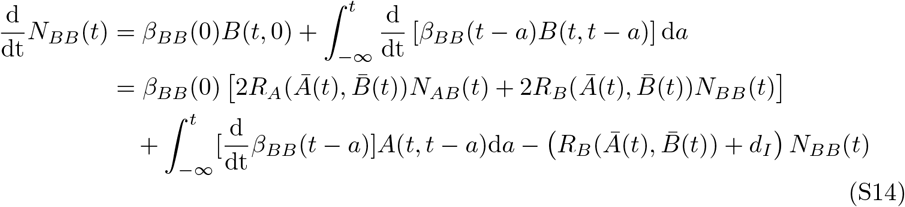

where we have used (S11) to write

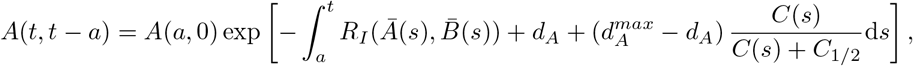

so that

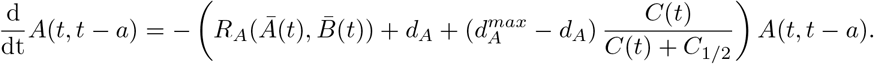

By using the following relationships for *β_AA_*(*t* — *a*) and *β_BB_*(*t* — *a*),

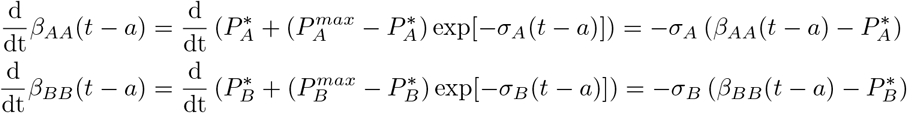

we simplify (S13) to

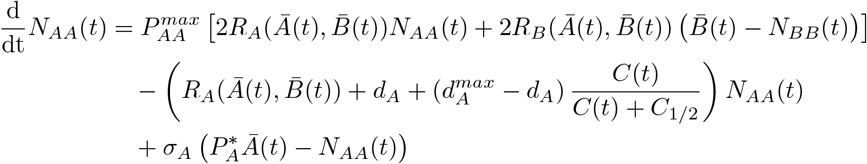

while (S14) becomes

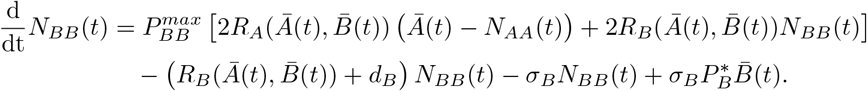

Thus, the system of ODEs for *Ā*(*t*) and 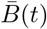 is

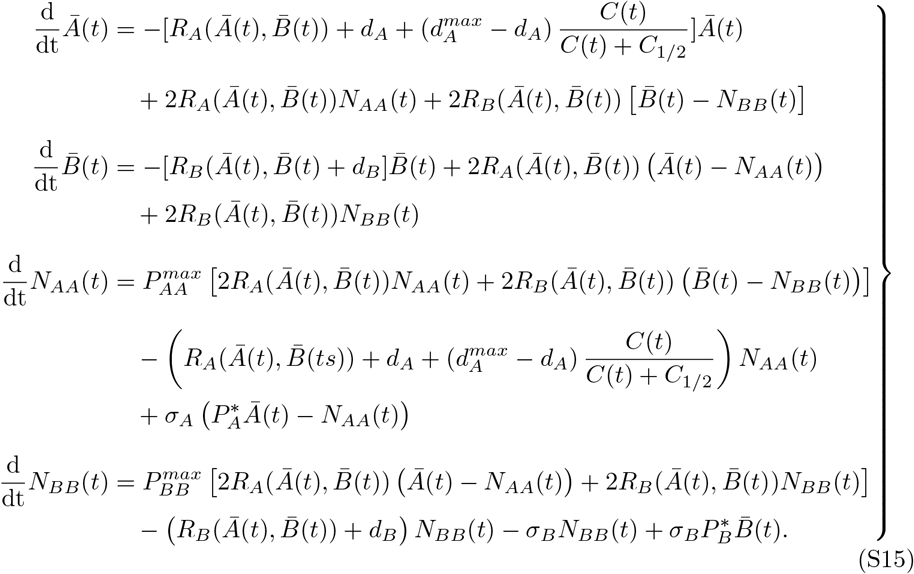

The ODE (S15) is intrinsically finite dimensional with real valued initial conditions given by 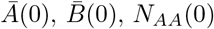 and *N_BB_*(0). This contrasts with the infinite dimensional system of PDEs (S2) with initial data given by *g_A_*(*a*) and *g_B_*(*a*) in the infinite dimensional space *L_1_*(0, ∞). To obtain the system of ODEs (S15), we partially solved the PDE (S2), so it is somewhat unsurprising that the resulting dynamical system is lower dimensional. We would like solutions of (S2) to correspond to solutions of (S15). Therefore, it is important to ensure that the initial conditions of the ODE system are appropriate. For integrable initial data *g_A_*(*a*) and *g_B_*(*a*) from the initial value problem (S2), it follows from the definition of *Ā*(*t*) and 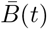 that

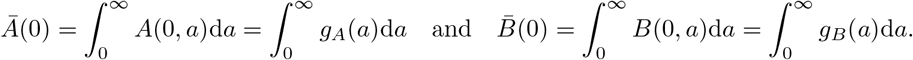

We can easily see that the initial conditions of *N_AA_* and *N_BB_* must satisfy

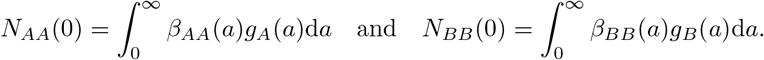

By the assumption that *g_A_*(*a*) and *g_B_*(*a*) are integrable and non-negative, these initial conditions are all finite and non-negative. In practice, it is simplest to assume that *g_A_*(*a*) and *g_B_* (*a*) are exponentially decaying functions of age so that the above integrals are simple to compute. This assumption of exponential decay in age is not unreasonable, both biologically and given the form of (S2).

#### Generic model parametrization

To study the role of phenotype switching on treatment resistance, we use a variety of physiologically based parameters rather than fitting the model to specific data. We assume that phenotype *A* cells successfully reproduce approximately once per day –similar to the reproductive time of most cells. Thus, we take *r_A_* = 0.7 ≈ log(2)/*t_A,2_·* Further, we assume that the phenotype *B* cells reproduce at about half the rate of phenotype *A* cells to account for the fitness cost of resistance [23,73], and set *r_B_* = 0.35. Unless otherwise stated, we fix *d_A_* = *d_B_* = 0. 01. We note that, with these parameters, drug sensitive cells are fitter than drug tolerant cells. We set 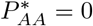 and 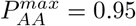. While this parametrization is deliberately generic, we show later that our results are robust to parameter variation.

### Model analysis

The mathematical model (S2) is quite simple and describes the time evolution of cell densities. In the following analysis, we do not consider the model with treatment, so *C*(*t*) = 0. We begin by demonstrating that solutions of (S2) evolving from non-negative and integrable initial data remain non-negative, as we would expect for a biological model.

#### Proposition S1.

*Let the model parameters be positive. Assume that g_A_(a) and g_B_(a) are integrable and almost-everywhere non-negative for a* ∈ (0, ∞). *Then, the solution of* (S2) *is non-negative for all time t* > 0 *and all growth functions* 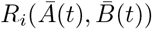.

*Proof*. Using the method of characteristics, the formal solution of (S2) is

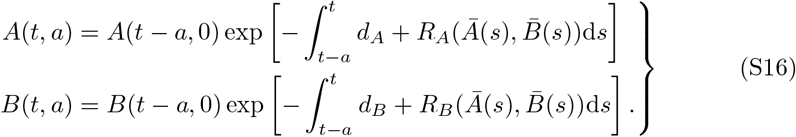

We note that *A*(*t, a*) and *B*(*t, a*) preserve the sign of *A*(*t* — *a*, 0) and *B*(*t* — *a*, 0) respectively, and we recall that *A*(*t* — *a*, 0) = *g_A_*(*a* — *t*) exp(—[*r_B_* + *d_B_*](*t* — *a*)) and *B*(*t* — *a*, 0) = *g_B_*(*a* — *t*) exp(—[*r_B_* + *d_B_*](*t* — *a*)) for *t* < *a*. Thus, to show that *A*(*t, a*) ⩾ 0 and *B*(*t, a*) ⩾ 0, it is sufficient to show that *A*(*x*, 0) ⩾ 0 and *B*(*x*, 0) ⩾ 0 for all *x* > 0.

We consider *A*(*x*, 0), as the same argument holds for *B*(*x*, 0), and proceed by contradiction. Assume for contradiction that *x*\* is the first time such that *A*(*x*\*, 0) < 0 or *B*(*x*\*, 0) < 0, so *A*(*s*, 0) ⩾ 0 and *B*(*s*, 0) ⩾ 0 for all *s* < *x*\*. Now, we must have

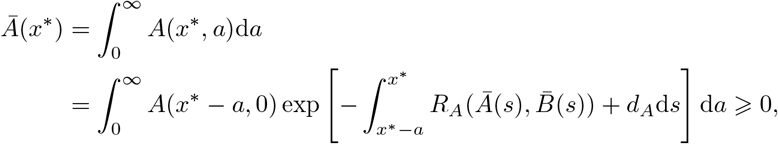

and

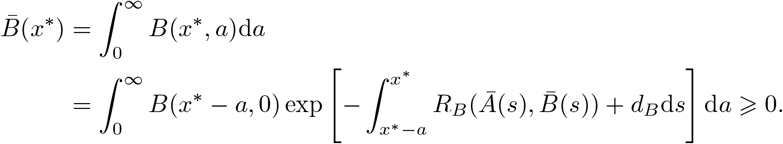

Therefore, the functions 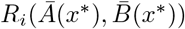 are non-negative for *i* = *A, B*. Furthermore, *β_ij_*(*0*) ⩾ 0 from definition. Then, using the definition of *A*(*x*, 0) given in (S3) and the formal solution of (S2), we calculate

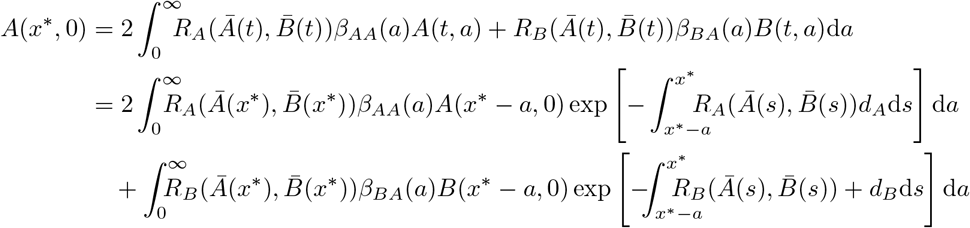

Finally, since *x*\* — *a* < *x*\*, it follows that *A*(*x*\* — *a*, 0) ⩾ 0 and *B*(*x*\* — *a*, 0) ⩾ 0, so the integrals

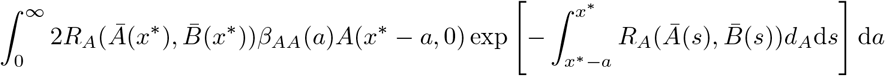

and

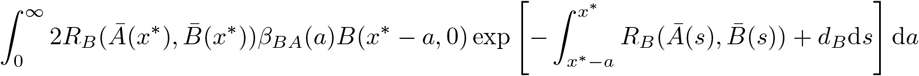

are integrals of non-negative functions over a set of positive measure. Thus, *A*(*x*\*, 0) is the sum of two non-negative integrals and must satisfy *A*(*x*\*, 0) ⩾ 0, a contradiction. The same argument for *B*(*x*\*, 0) yields the claim.

Turning now to the ODE model for *Ā*(*t*) and 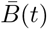 and having prescribed appropriate initial data, the theory of ODEs ensures that the initial value problem (IVP) defined by (S15) has a unique solution. Further, it follows immediately from Proposition S1 that solutions of (S15) evolving from non-negative initial data remain non-negative.

#### Nonlinear Eigenproblem for the Malthusian Parameter

To analyse the long term behaviour of the cell population, we search for an stable age distribution in the population [63, 69]. This stable age distribution is equivalent to finding the first eigenelements of (S2). We assume a solution of the type

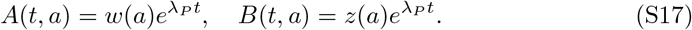

where λ*_P_* is the Malthusian parameter to be determined [63]. The Malthusian parameter is an important quantity in population dynamics [47, 66, 74], and is typically used as measure of population fitness [41, 75]. Later, we will show that the expected sign relationship between the Malthusian parameter λ*_P_* and *R*_0_ — 1, where the basic reproduction number *R*_0_ is another classical measure of population fitness, holds in our model. This result normally follows immediately in most structured population models. However, the inclusion of phenotypic switching in our model complicates this relationship.

The unknown functions *w*(*a*) and *z*(*a*) are the age distributions of *A* and *B* respectively. These functions define a system of ordinary differential equations (ODEs) from which we will obtain a nonlinear eigenvalue problem with solution λ*_P_*. For general λ, inserting the ansatz (S17) into (S2) yields the following system of ODEs

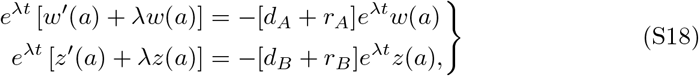

with solutions

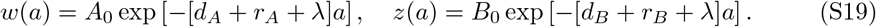

We must now ensure agreement with the boundary condition (S3) of the population PDE (S2), so

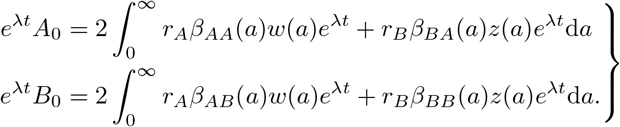

Cancelling the e^λt^ terms gives a system of equations for the unknowns λ, *A_0_* and *B*_0_

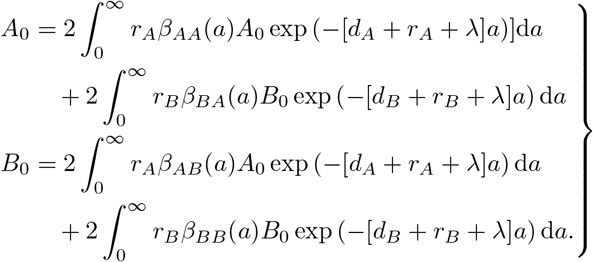

This linear system for the unknowns *A*_0_, *B*_0_ is equivalent to

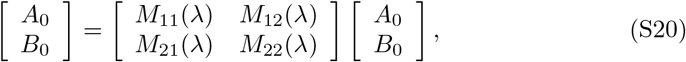

where

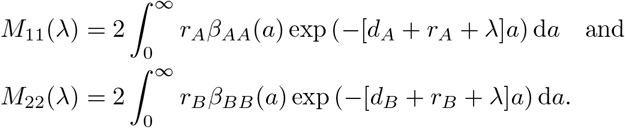

We note that newly born cells must be of either phenotype *A* or *B*, so we have

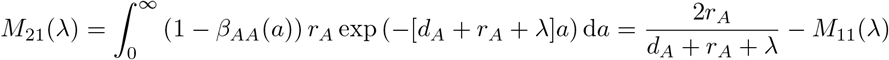

and

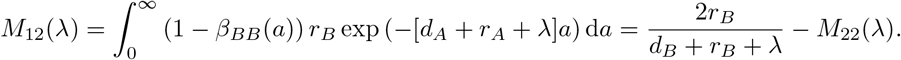

Consequently, the Malthusian parameter λ*_P_* is the rightmost real solution of the nonlinear eigenproblem defined by (S20) and must satisfy

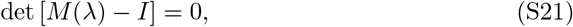

where *M*(λ) is given by the matrix in (S20). In the case of no phenotypic switching, the matrix *M*(λ) is diagonal and this eigenvalue problem is simple. The following proposition is nearly obvious from the biological interpretation of the problem.

##### Proposition S2

*Assume that cells cannot switch phenotype and that the model parameters are positive. Then, the Malthusian parameter is given by* λ*_P_* = max[*r_A_* — *d_A_*, *r_B_* — *r_B_*].

*Proof*. If offspring directly inherit the phenotype of the parent cell, then 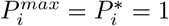. It follows that *M*_12_ = *M*_21_ = 0, and the matrix (S20) is given by

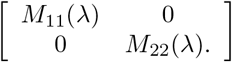

The eigenvalues are therefore *M*_11_(λ) and *M*_22_(λ), which, for 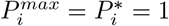, are given by

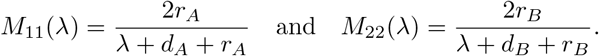

Then, 1 is an eigenvalue if and only if λ = *r_A_* — *d_A_* or λ = *r_B_* — *d_B_*. The Malthusian parameter is the maximum of these values, so λ*_P_* = max[*r_A_* — *d_A_*, *r_B_* — *d_B_*].

It follows from the preceding proposition that, if no phenotypic switching can occur, a population comprised of entirely phenotype *A* cells has Malthusian parameter λ*_A_* = *r_A_* — *d_A_* and a population with only type *B* cells has Malthusian parameter λ*_B_* = *γ_B_* — *d_B_*. To simplify notation, we will assume, without loss of generality, that λ*_B_* ⩽ λ*_A_*.

We will now show that allowing for phenotypic switching acts to decrease population fitness. Namely, if λ*_P_* is the Malthusian parameter of the switching population, then λ*_P_* ∈ (λ*_B_*, λ*_A_*). We begin by removing the restriction that 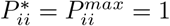, and evaluate (S21). Then, this determinant becomes

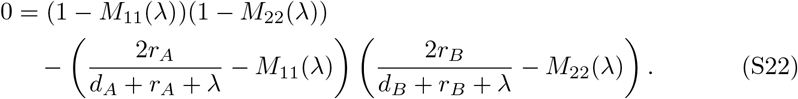

For *β_ii_* given by (S4) and (S5), we calculate

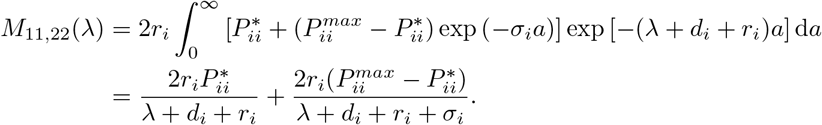

Then, equation (S22) becomes

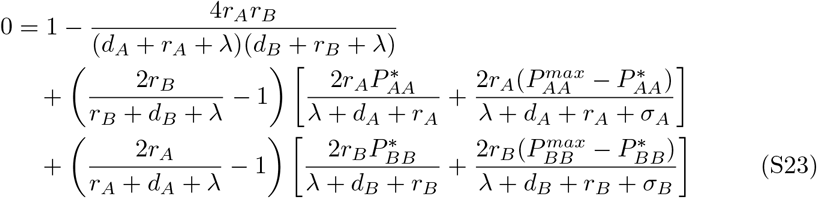

To simplify notation in the following analysis, we set

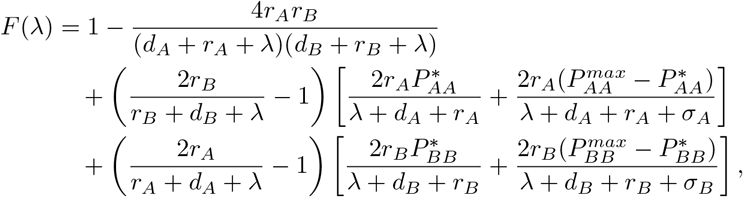

so that roots of *F*(λ) correspond to solutions of (S23).

It remains to show that (S23) admits at least one real root. While *F*(λ) may admit multiple real roots, the Malthusian parameter λ*_P_* is the rightmost real root by convention. As we have seen, in models without phenotypic switching, it is typically fairly straightforward to prove that λ* exists and is unique [63,69]. However, the phenotypic switching in (S2) results in the off diagonal terms of *M*(λ) and complicates the analysis here, as the Malthusian parameter is no longer a strictly monotonic function of the parameters *r_i_, d_i_*. However, *F*(λ) is eventually monotonic for λ > 0 large enough.

##### Lemma S3

*F*(λ) *is strictly increasing for* λ > λ*_A_* = max(λ*_A_, λ_B_*).

*Proof*. To simplify notation in the proof, we write for *i* = *A, B*,

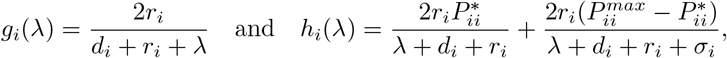

so

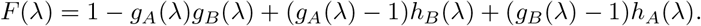

Then, differentiating *F*(λ) and regrouping terms gives

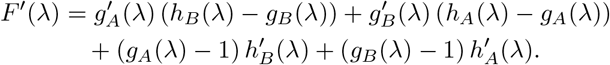

It is clear that 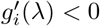 and 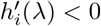. Further, for λ > max(*r_A_, — d_A_, r_B_*, — *d_B_*), we obtain *g_i_*(λ) — 1 < 0 and

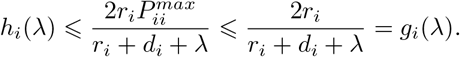

Therefore, each of the terms in *F′*(λ) is the product of two non-positive functions, with

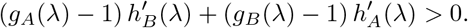

It follows that *F′*(λ) > 0 and *F*(λ) is strictly increasing for λ > max(*r_A_* — *d_A_, r_B_* — *d_B_*)

We continue by considering an extremely particular case, where both sub-populations have the same fitness. Consequently, phenotypic switching does not affect population fitness.

##### Lemma S4

*Let the model parameters be positive. If r_A_* — *d_A_* = *r_B_* — *d_B_, then* λ*_P_* = *r_A_* — *d_A_* = *r_B_* — *d_B_ is the Malthusian parameter*.

*Proof*. Evaluating (S22) at λ* = *r_A_* — *d_A_* = *r_B_* — *d_B_* gives

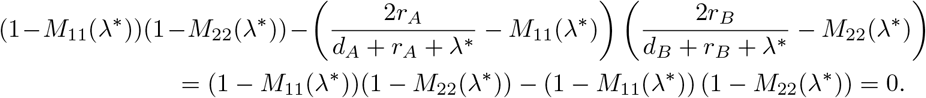

Then, since *F*(λ) is strictly increasing for λ > *r_A_* — *d_A_*, λ* is the rightmost real root of *F*, and λ*_P_* = λ* = *r_A_* — *d_A_*.

We now consider the more general case, where *r_A_* — *d_A_* ≠ *r_B_* — *d_B_*.

##### Lemma S5

*Let the model parameters be positive and assume that* λ*_A_* > λ*_B_* > — min[*r_A_* + *d_A_,r_B_* + *d_B_*]. *Then, there exists a real root* λ* *of* (S23) *with* λ* ∈ (λ*_B_*, ∞).

*Proof*. We begin by noting *F*(λ) is continuous and well-defined for λ ∈ (— min[*r_A_* + *d_A_,r_B_* + *d_B_*], ∞). Further,

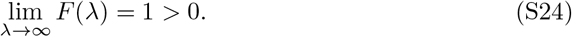

Now, recall that λ*_B_* = *r_B_* — *d_B_*, so *r_B_* + *d_B_* + λ*_B_* = 2r*_B_* and calculate

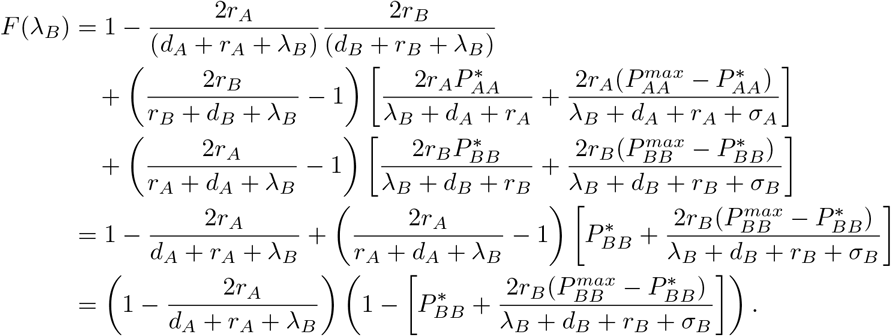

Now, since σ*_B_* > 0, λ*_B_* + *d_B_* + *r_B_* + σ*_B_* = σ*_B_* + 2*r_B_* > 2*r_B_* and

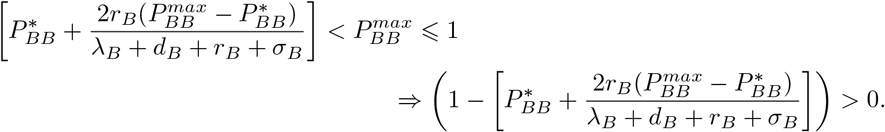

By the assumption that λ*_B_* < λ*_A_*, we have *r_A_* + *d_A_* + λ*_B_* < 2*_A_*. It follows that

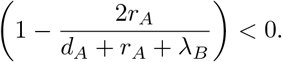

Thus, *F*(λ*_B_*) < 0 and the intermediate value theorem yields the claim.

We now regroup Lemmas S3 and S5 to establish the existence of the Malthusian parameter.

##### Theorem S6

*Let the model parameters be positive. Then the Malthusian parameter* λ*_P_ of the population with phenotypic switching satisfies* λ*_P_* ∈ (λ*_B_*, λ*_A_*).

*Proof*. The existence and lower bound follows immediately from Lemma S3 and Lemma S5. It only remains to show that λ*_P_* < λ*_A_*. Recalling that λ*_A_* = *r_A_* — *d_A_*, we calculate

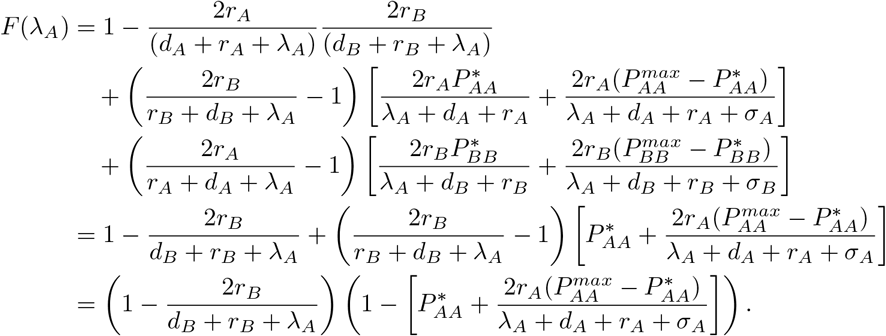

From the definition of λ*_A_*, it follows that λ*_A_* + *d_A_* + *r_A_* + σ*_A_* > 2*r_A_*, so

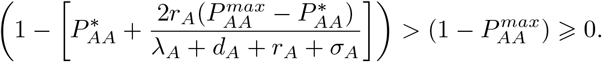

Moreover, λ*_A_* + *r_B_* + *d_B_* = *r_A_* — *d_A_* + *r_B_* + *d_B_* > *r_B_* — *d_B_* + *r_B_* + *d_B_* = 2*r_B_*, so

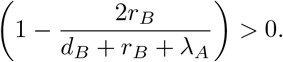

Thus, *F*(λ*_A_*) is the product of two positive terms which ensures that *F*(λ*_A_*) > 0. The intermediate value theorem ensures that there is at least one root 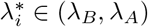. Finally, Lemma S3 ensures that *F*(λ) has no real roots λ > *r_A_* — *d_A_*. Then, the Malthusian parameter is the maximum of the possible roots 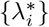 in the interval (λ*_B_*, λ*_A_*), so 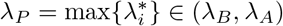.

#### The Basic Reproduction Number

As previously mentioned, there typically is a correspondence between the Malthusian parameter λ*_P_* and the basic reproduction number. The basic reproduction number is the spectral radius of the next generation operator [46], and is typically understood as the expected number of new cells produced by each existing cell. We now recast the nonlinear eigenproblem (S20) as a renewal type equation from which we derive the basic reproduction number. In what follows, we use the formal solution of (S2) given by (S16).

To simplify notation, define

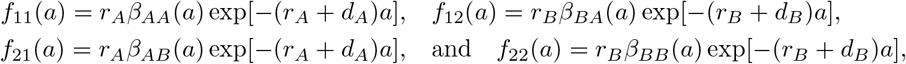

and note that the boundary terms *A*(*t, 0*) and *B*(*t, 0*) are functions of time. We recall that we extended the initial conditions *g_A_*(*a*) and *g_B_*(*a*) to define *A*(*t* — *a*, 0) and *B*(*t* — *a*, 0) for *t* < *a* by *A*(*t* — *a*, 0) = *g_A_*(*a* — *t*) exp(—[*r_B_* + *d_B_*](*t* — *a*)) and *B*(*t* — *a*, 0) = *g_B_*(*a* — *t*) exp(—[*r_B_* + *d_B_*](*t* — *a*)).

Now, inserting the formal solution (S16) into the boundary condition (S3), we see that *A*(*t, 0*) and *B*(*t, 0*) satisfy the renewal equation

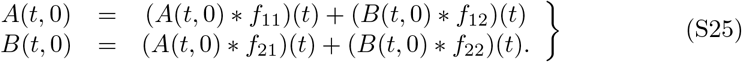

Taking the Laplace transform of (S25) gives the linear system

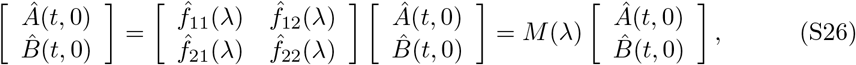

where *M*(λ) is given by (S20). The untreated next generation operator (NGO) is therefore given by [46, 76, 77]

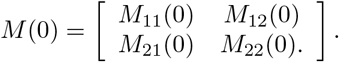

and *R*_0_ is the spectral radius of *M*(0). The eigenvalues of *M*(0) are

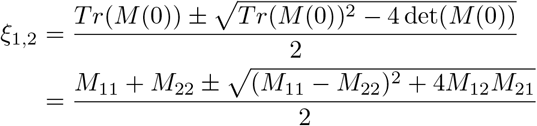

In particular, we note that ξ _12_ are real numbers, and the reproductive number of the mixed population is

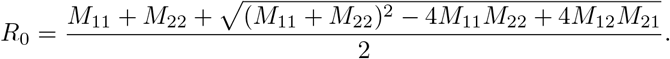

As each individual cell can only produce a maximum of two daughter cells, we expect *R*_0_ ⩽ 2. We now show that this is the case, and that this bound will be reached only if there is no death.

##### Lemma S7

*Let the model parameters be non-negative. Then*, 0 ⩽ *R*_0_ ⩽ 2, *and achieves these bounds if r_A_* = *r_B_* = 0, *or if d_A_* = *d_B_* =0 *respectively*.

*Proof*. The matrix *M*(λ) is comprised of non-negative elements *M_ij_*(λ), so

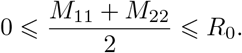

Now, *R*_0_ = 0 if and only if

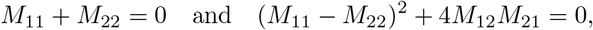

which can only be achieved when *r_A_* = *r_B_* = 0. To show the upper bound, we recall that

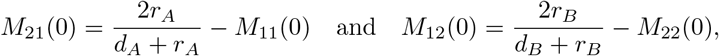

so the NGO is given by

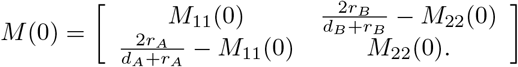

Then, the Gershgorin circle theorem implies that

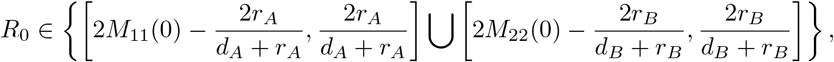

so

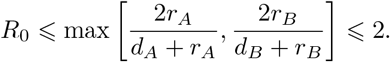

with strict inequality if both *d_A_* > 0 and *d_B_* > 0. Now, assume that *d_A_* = *d_B_* = 0, so that *M*(0)*^T^* becomes

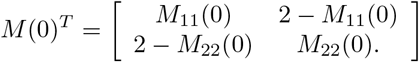

which has spectral radius 2. Since *M*(0) and *M*(0)*^T^* are similar, it follows that *R*_0_ = 2 if *d_A_* = *d_B_* = 0.

Similar to the Malthusian parameter, the basic reproduction number is a measure of population fitness, where the sign of *R*_0_ — 1 determines if cells are expected to replace themselves through replication. When the birth and death rates are balanced, we now show that we should not expect population growth.

##### Lemma S8

*Let the model parameters be positive and assume that r_A_* — *d_A_* = *r_B_* — *d_B_* = 0. *Then* λ*_P_* = *R*_0_ — 1 = 0.

*Proof*. Since *r_A_* = *d_A_* and *r_B_* = *d_B_*, it follows that

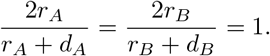

Once again using the similarity between *M*(0) and *M*(0)*^T^*, we compute

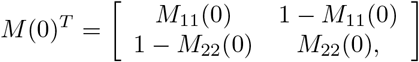

which clearly has spectral radius 1, so the *R*_0_ claim in shown. Further, for *F*(λ) given by (S23), it is simple to see that *F*(0) = 0 when λ*_A_* = λ*_B_* = 0. Further, Lemma S3 shows that *F*(λ) is strictly increasing for λ > max(*r_A_* — *d_A_, r_B_* — *d_B_*) = 0. Therefore, 0 is the rightmost real root of *F*(λ) and λ*_P_* = 0.

##### Theorem S9

*If the model parameters are positive, then, sign*(λ*_P_*) = *sign*(*R*_0_ — 1).

*Proof*. Consider the spectral radius of *M*(λ) given by

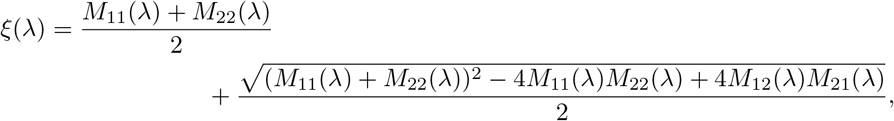

and note that *R*_0_ = *ξ*(0) while the Malthusian parameter λ*_P_* satisfies *ξ*(λ*_P_*) = 1. Viewing *ξ* as a function of *M_ij_* for *i, j* = 1, 2, we compute

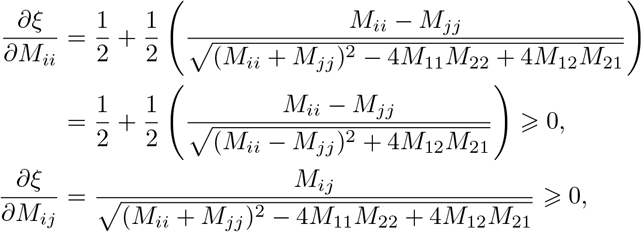

where we have used the non-negativity of *M_ij_* to establish the sign of 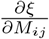. Since each *M_ij_* is strictly decreasing in λ, it follows that

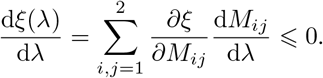

If one of *M_ij_* ≠ 0, then the above inequality is strict, and *ξ* is a strictly decreasing function of λ. Moreover, *M_ij_* = 0 for all *i, j* can only occur when *r_A_* = *r_B_* = 0. Therefore, for positive model parameters, *ξ*(λ) is a strictly decreasing function.

Now, assume that λ*_P_* > 0, so *R*_0_ = *ξ*(0) > *ξ*(λ*_P_*) = 1. Conversely, assume that *R*_0_ — 1 > 0, so then *ξ*(0) > 1. As

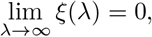

the intermediate value theorem ensures that λ_P_ > 0. Next, assume that *R*_0_ = *ξ*(0) < 1. Then, as λ*_P_* must exist from Theorem S6, the monotonicity of *ξ* gives λ*_P_* < 0. Finally, assume that λ*_P_* < 0. It follows that 1 = *ξ*(λ*_P_*) > *ξ*(0) = *R*_0_.

The sign relationship between the Malthusian paramether λ*_P_* and the basic reproduction number *R*_0_ established in Theorem S9 allows us to determine if the tumour population will grow or decay through a number of techniques. As we will show later, it is sufficient to design a treatment schedule to ensure that *R*_0_ < 1, and the sign relationship established in Theorem S9 immediately yields that λ*_P_* < 0 so small tumour population cannot grow. Conversely, if both λ*_A_* < 0 and λ*_B_* < 0, Theorem S6 implies that λ*_P_* < 0, so *R*_0_ < 1 from which it follows that the tumour population is decaying.

#### Stable Age Distribution and Population Proportion

Having calculated the Malthusian parameter, we can determine the stable age distribution. From the non-linear eigenproblem (S20), each value of the Malthusian parameter λ*_P_* defines an eigenvector [*A_0_*, *B_0_*] and corresponding solution to (S18) given by (S19). At a given time *t*, these exponential functions model the proportion of cells born at time *t* — *a* that have not died or reproduced. Perhaps counter-intuitively, a larger value of λ implies that there are fewer cells for a given age a. However, as nascent cells are necessarily born with age 0, an accumulation of old cells (large a) is indicative of a population that is not reproducing.

Finally, after solving the non-linear eigenproblem for the Malthusian parameter, we can calculate the explicit solution of (S2) under steady state growth using (S17). Then, it is simple to calculate

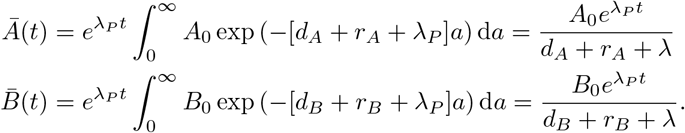

In practice, it is difficult to calculate the age of each cell in a cohort, but relatively easy to calculate the proportion of different phenotypes. Thus, the ratio

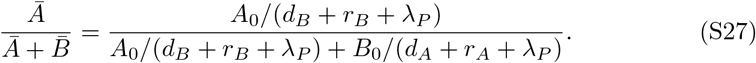

is likely to be of clinical interest in understanding drug resistance, although the assumption of Malthusian growth is only appropriate in the case of unlimited resources. We note that this ratio is dependent on both the Malthusian parameter λ*_P_* and the eigendirection corresponding to λ*_P_* through [*A_0_, B_0_*]. Later, we will show how this ratio is dependent on the model parameters and growth function 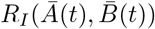. This dependence on the growth function indicates that the ratio 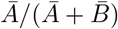 will evolve as a population exhausts available resources. The importance of growth phase on population make up is well established in *E.coli* populations that exhibit bet-hedging [13, 20, 34].

#### Frequency dependent fitness outperforms Malthusian growth

The ability to calculate the Malthusian parameter relies on the existence of a dominant exponential type solution. Exponential growth is unrealistic in the case of finite resources, as growing populations will exhaust available resources. Therefore, we also consider the dynamics of *N*(*t*) for logistic and Allee type growth functions, given by (S7) and (S8) respectively. To study growth in resource rich environments, we take *N*(*0*) ≪ *K*, so there are sufficient resources available initially.

In populations dominated by the “fitter” drug sensitive phenotype, it is reasonable to expect Malthusian growth to dominate resource limited growth, even in the case *N*(*0*) ≪ K. Biologically, this corresponds to the competition for finitely many resources limiting growth, even for small populations. However, we show in Figure S2 that it is possible that cooperation amongst drug tolerant cells can initially out perform exponential type growth.

In Figure S2, we plot the ratio of the resource limited vs unlimited growth, so Malthusian growth would correspond to a horizontal line at 1. As shown in Figure S2 (a), a majority drug sensitive initial population can briefly match, or even slightly surpass (due to a slight Allee effect), Malthusian growth. As the Malthusian parameter falls between the fitnesses of phenotype *A* and *B*, a population initially comprised of exclusively drug sensitive cells will outperform Malthusian growth of the mixed population until the effects of phenotypic switching become apparent and the drug tolerant population grows in size. Moreover, as *N*(*t*) increases and cells compete for limited resources, Malthusian growth overtakes and dominates the finite resource case. As drug sensitive cells are assumed to be fitter than drug tolerant cells, the presence of less fit drug tolerant cells both consumes resources and also lowers the average fitness of the population.

**Fig S2.**
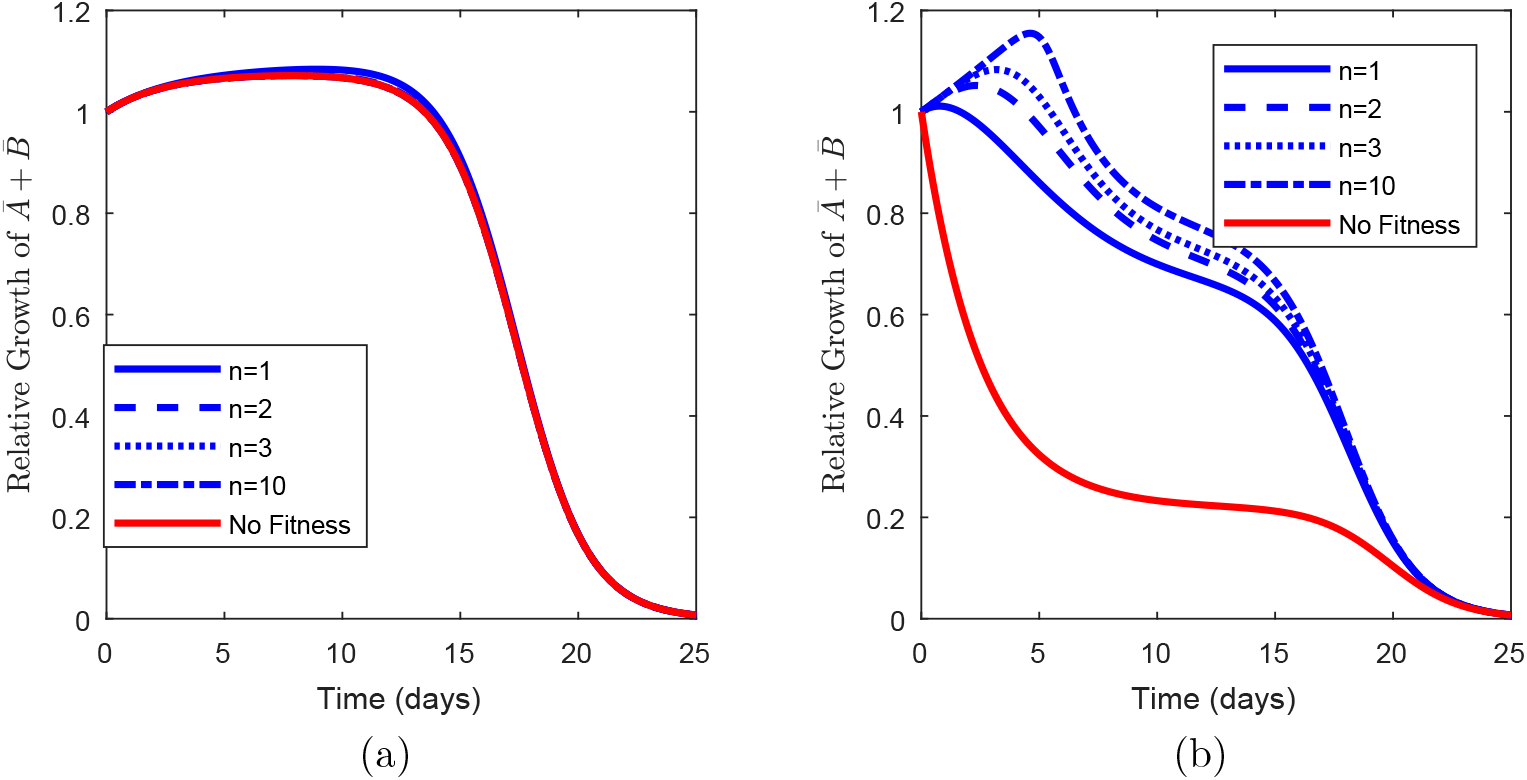
A comparison of growth rates for different growth functions *f_n_, n* = 1, 2, 3, 10, against Malthusian growth. The “no fitness” curves corresponds to no frequency dependent fitness increase and *f_n_* = 1. Figure (a) shows population evolution from an initial population comprised of 100 drug sensitive cells and one drug tolerant cell. Conversely, Figure (b) shows population evolution from an initial population comprised of one phenotype *B* cell, or 1 drug sensitive cell and 100 drug tolerant cells.

Conversely, Figure S2 (b) presents a much more interesting situation. From an initial population of one drug sensitive cell and 100 drug tolerant cells, the impact of the Allee effect drastically changes initial population growth. From this high initial proportion of drug tolerant cells, the constrained growth model out performs Malthusian growth. Since the drug tolerant cells receive a substantial increase in fitness due to the Allee effect, the initial fitness of the population is higher – despite the finite amount of resources – than the fitness of the total population in the presence of unlimited resources and no cooperation. However, despite the cooperation induced increased fitness of drug tolerant cells, the population eventually evolves towards a predominantly drug sensitive population due to phenotypic switching and the growth rate falls below Malthusian growth.

#### Parameter identifiability during cancer therapy

Thus far, we have shown that the phenotypic switching strategy employed by a population can lead to different types of therapeutic resistance. Here, we discuss the difficulties of determining the switching probability based on untreated population data. We consider *in vitro* data from the growth of multiple myeloma growth in mice [78], and numerous different phenotypic switching strategies. After digitizing the data from Figure 1 (a) of [78] and fixing a phenotypic switching strategy, we fit the tumour growth parameters *r_A_,r_B_* and *d_A_* = *d_B_* to the time series by minimizing

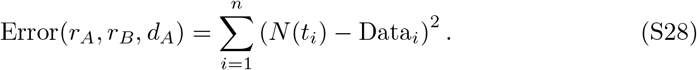

The data from [78] is sampled at times 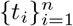. For a given phenotypic switching strategy and parameter set *r_A_, r_B_* and *d_A_* = *d_B_*, we simulate (S15) and sample the numerical solution at the times *t_i_*. Equation (S28) is then the *l_2_* distance between *N*(*t_i_*) and the [78] data. In addition to the switch and stay strategies discussed in the Main Text, we consider 6 additional phenotypic strategies given by the pairs 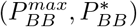: (1,0.25); (1,0.5); (1,0.75); (0.9,0.25); (0.9, 0.5); and (0.9, 0.75).

**Fig S3.**
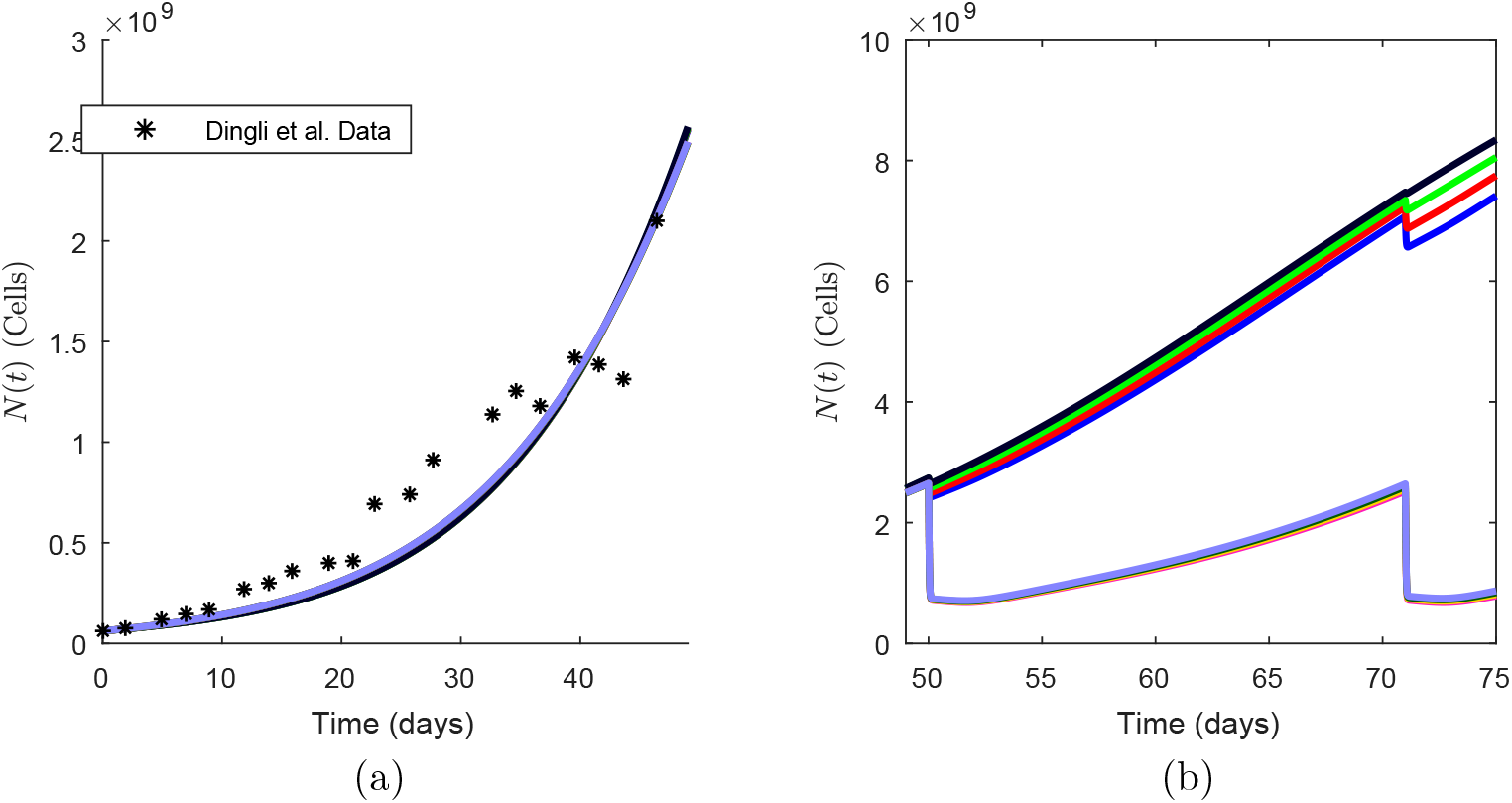
Fitting of the mathematical model to the Dingli 2009 data for a variety of fitting strategies. Figure (a) shows the fitting of the mathematical model (S2) to the Dingli 2009 data for 8 different switching strategies. Figure (b) shows the same 8 strategies after 2 applications of therapy.

In Figure S3 (a), we show the fitting results for each of the 8 different switching strategies to the [78] data. The eight curves are essentially indistinguishable, which indicates a possible difficulty in translating our results to the clinic. In Figure S3 (b), we simulate two therapy cycles, and the eight previously indistinguishable curves are then separated into two clusters almost immediately following therapy. The four strategies corresponding to the less responsive (larger *N*(*t*): black, green, red and blue) curves all have 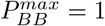, while the four strategies corresponding to higher response to therapy (lower *N*(*t*): purple and other colors) all have 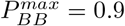. Thus, we see that population response to treatment can stratify populations by their switching strategy. Therefore, it should be possible to use treatment response to determine the approximate switching strategy of a tumour biopsy and use this information to inform therapeutic strategies.

#### Generic strategy to avoid treatment failure

For the generic tumour growth parameters used thus far, *r_B_/r_A_* = 1/2. While our results are robust to different values of *ε*, we illustrate our results with *ε* = 0.7 in the threshold ratio 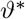 from the Main Text Eq. (5). To test if this simple threshold ratio is sufficient to avoid the establishment of therapy, we follow the same periodic dosing as shown in Main Text Fig 3, but we only administer therapy if the ratio 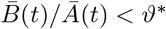 and label this strategy *adjustable therapy*.

However, it is unrealistic that clinicians will determine the ratio 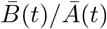 and immediately administer therapy. Therefore, to decide if treatment will be applied at time *t*\*, we consider 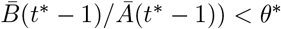 which corresponds to clinicians taking one day to complete the phenotype profile of the tumour. We show that adjustable therapy avoids the establishment of resistance in Figure S4. As in the Zhang et al. [22] trial, the main benefit of this adjustable therapy is, that by avoiding the development of resistance, therapy with the same drug can continue indefinitely. In particular, the effectiveness of the *adjustable therapy*, as measured by disease burden (S29), increases the longer that therapy is applied. We note that, rather than using the ratio of drug tolerant to sensitive cells to determine if therapy should be applied, it is possible to use a cancer specific biomarker [22].

Inspired from the success of adaptive therapy in prostate cancer [5, 22], we developed a simple strategy to avoid the establishment of resistance in *Avoiding the establishment of a drug tolerant population*. We show the results from that section here. It is important to note, once again, that our model is quite coarse, so these results serve more as a proof-of-concept, rather than a proposed therapeutic strategy. As such, and to avoid confusion, we refer to the following strategy as *adjustable therapy*. We measure treatment efficiency by

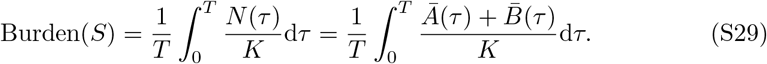

**Fig S4.**
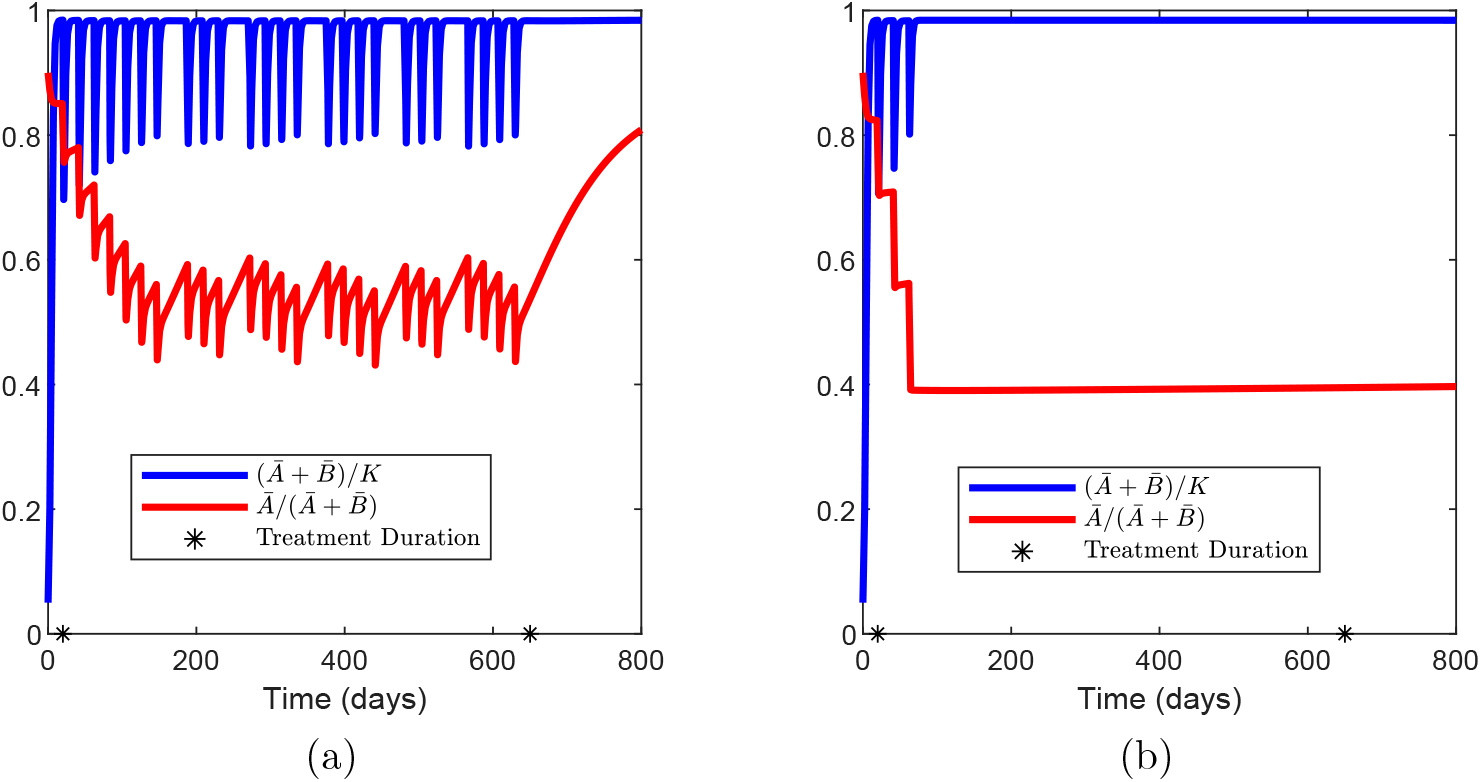
The effect of adjustable therapy on a population using either a *switch* (panel (a)) or *stay* (panel (b)) strategy. In both cases, treatment is applied between the black stars, while the red curve shows the proportion 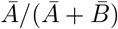, and the blue curve shows the dynamics of *N*(*t*).

Figure S4 demonstrates that the proposed adjustable therapy strategies avoid the establishment of a dominant resistant phenotype. We compare the results of the adaptive therapy against periodic treatment in Figure S4. Figure S4 a shows the response of a population with a switch strategy 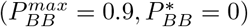 and transient resistance to adjustable therapy, while Figure S4 b shows the same effect of adjustable therapy on a population with a stay strategy and permanent resistance 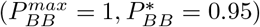. When comparing the effectiveness of the therapy using (S29) over 240 days of treatment, the adjustable therapy is leads to a disease burden that is 0.2% lower (for the switch strategy) or 0.03% higher (for the stay strategy) than the periodic therapy in Figure S4 a and b, respectively.

If these adjustable strategies are continued for longer than 240 days, there is an increasingly important decrease in disease burden when compared to the periodic type treatment resulting from the eventual ineffectiveness of the periodic dosing due to therapeutic resistance that occurs when the population is dominated by the resistant phenotype.

#### General results are robust to parameter variation

We have thus far shown that changes in phenotypic switching strategy can lead to two distinct types of therapy resistance for a given parameter set. Here, we confirm that this qualitative result is robust to parameter variations, and that the results presented in the main text are representative of model dynamics for a number of different parametrizations. We continue to enforce *d_A_* = *d_B_* and explore the possible parameter combinations by selecting triplets of parameter values from the set

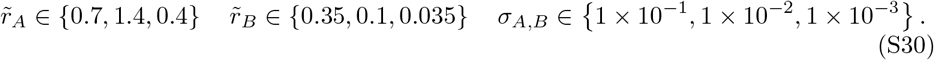

For a given parameter combination, we simulated the mathematical model with periodic treatment as described previously.

Each of the 27 possible parameter combinations were tested with both the switch and stay strategies for the drug tolerant population and each demonstrated transient resistance and the re-establishment of the wild type phenotype shortly after cessation of therapy when coupled with a switch strategy, similar to Main Text Fig 3 (a). Conversely, when simulated with a stay strategy, each parameter combination demonstrated the permanent resistance shown in Main Text Fig 3 (b). Therefore, we consider the simulations shown to be representative.

As demonstrated, the switch and stay strategies are crucial in determining the appearance and duration of therapeutic resistance. However, the probabilities 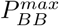 and 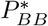 used to determine the switch and stay strategies are vastly different. Therefore, we test the robustness of the qualitative results shown in Main Text Fig 3 for different extremes of the switch and stay strategies. Once again, we consider two distinct strategies. In the first, we fix 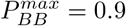, as in the switch strategy, and test different values of 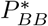 from 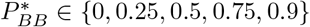. Then, for each value of 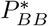, we simulate periodic therapy. For 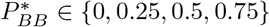, resistance was transient and the wild type phenotype was re-established in the population shortly after the end of therapy. Conversely, for 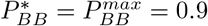, re-establishment of the wild type population took over 800 days after treatment cessation, which is effectively permanent resistance. We note that, if 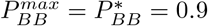, then reproductive resistant cells are quite likely to transfer their phenotype to offspring, and this strategy closely resembles a staying strategy. As a consequence, the correspondence between transient resistance and a “switch” strategy appears to be robust to parameter changes.

Now, to test if a stay strategy consistently predicts permanent resistance, we fix the homoeostatic switching probability 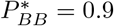 and vary 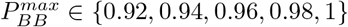. For all values of 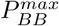, periodic therapy led to resistant phenotype dominance that persisted for over 800 days following the end of therapy. Once again, in the context of therapeutic scheduling, this resistance is effectively permanent. Thus, the stay strategy consistently leads to permanent treatment resistance, and we consider our results in the Main Text to be representative.

#### Effectiveness of Adjustable Therapy

We also verified the effectiveness of adjustable therapy for the combination of growth parameters given in (S30). Once again, we tested all 27 possible combinations of parameters in (S30) for both the switch and stay strategy, while holding the measure of acquired fitness, *ε*, fixed at *ε* = 0.7. For the switch strategy, adjustable therapy improved upon periodic therapy in 15/27 cases, with 8 of the remaining cases showing less than a 0.0001% increase in tumour burden. In the worst case, adjustable therapy led to an increase in tumour burden by 3%, while in the best case, there was a 2% decrease.

We completed the same test for a population with a stay strategy. There, adjustable therapy improved upon periodic therapy in 15/27 cases. In the worst case, adjustable therapy led to an increase in tumour burden by 0.6%, while in the best case, there was a 0.2% decrease. Therefore, the therapeutic improvement offered by adjustable therapy appears to be robust to parameter variations. Moreover, the adjustable schedule consistently avoided the establishment of resistance.

These results differ from the sustained treatment success in for NSCLC in the main text for an important reason, namely that, for the generic parametrization considered here, λ*_B_* > 0. Accordingly, our analysis and derivation of the model informed therapy in the main text does not apply. Therefore we are not using an optimized dose size, and as the drug tolerant cells are entirely resistant to therapy, it is not possible to drive 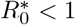.

### Application to Non-Small Cell Lung Cancer

Similar to the experiments that identified persisters in bacterial populations [26, 34], the experimental set up used by Craig et al. [7] begins with a constant environment and a genotypically homogeneous population of cancer cells. Thus, it may be tempting to conclude that the phenotypic heterogeneity present is solely due to stochastic phenotype switching. However, the distinction between phenotypic plasticity, wherein cells change phenotype in response to environmental change, and truly stochastic phenotype switching is subtle [26]. Moreover, this dichotomic representation of phenotypic heterogeneity does not account for partially heritable phenotype, as reported by Yang et al. [31] and considered in our model. Nevertheless, the NSCLC data offers an initial opportunity to apply our simple mathematical model to cancer data and explore the role of phenotype switching in treatment resistance.

#### Parameter fitting

Here, we present the results of the fitting procedure for the WT, M1 and M2 data from [7]. As mentioned in the Main Text, we fit this data by minimizing the *l_2_* error between the data and the model simulation. We used the algorithm *fmincon* [64] with 15 distinct initial points. We show the results of the fitting for the WT, M1 and M2 data in Figures S5 and S6, and give the parameter values in Table S1 and S2.

**Fig S5.**
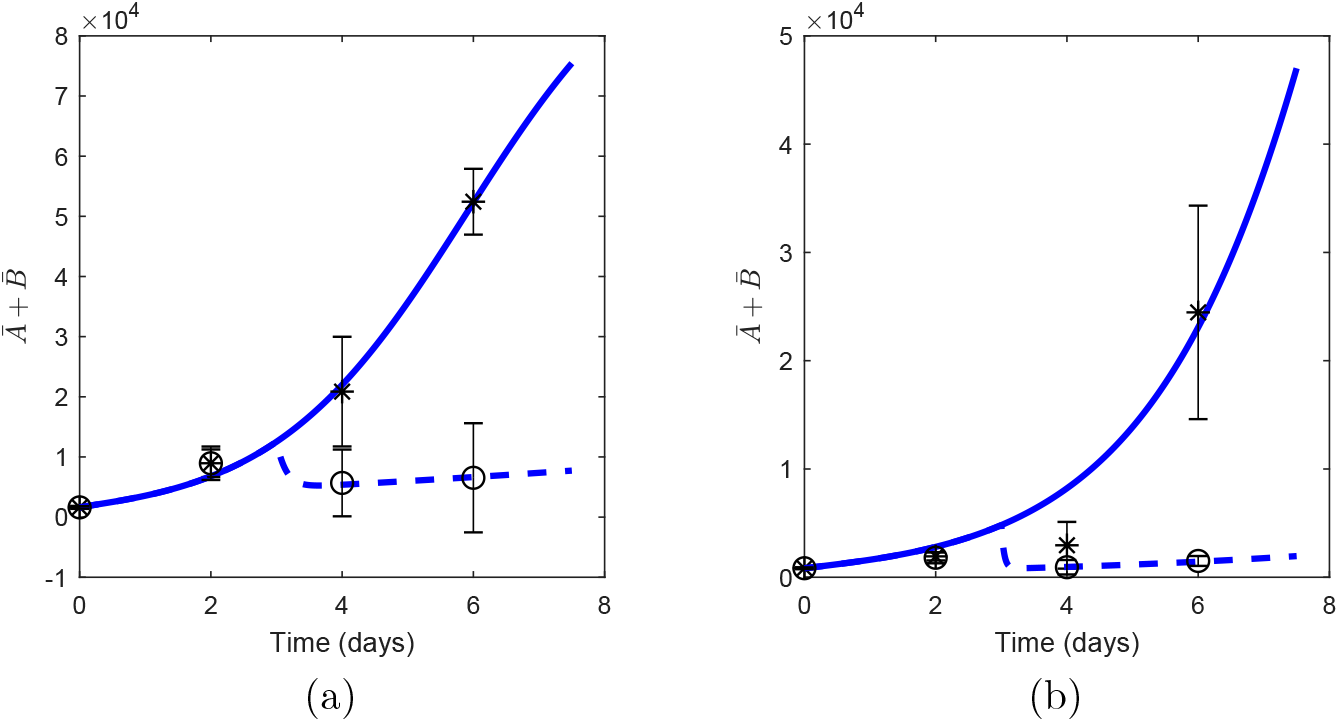
Fitting results of Equation (S2) to the WT and M1 data from [7] in Figures (a) and (b), respectively. In all cases, the untreated data is given by the black stars while the untreated simulation is in solid blue. The docetaxel treated data is given by the hollow circles and the treated simulation is in dashed blue.

**Fig S6.**
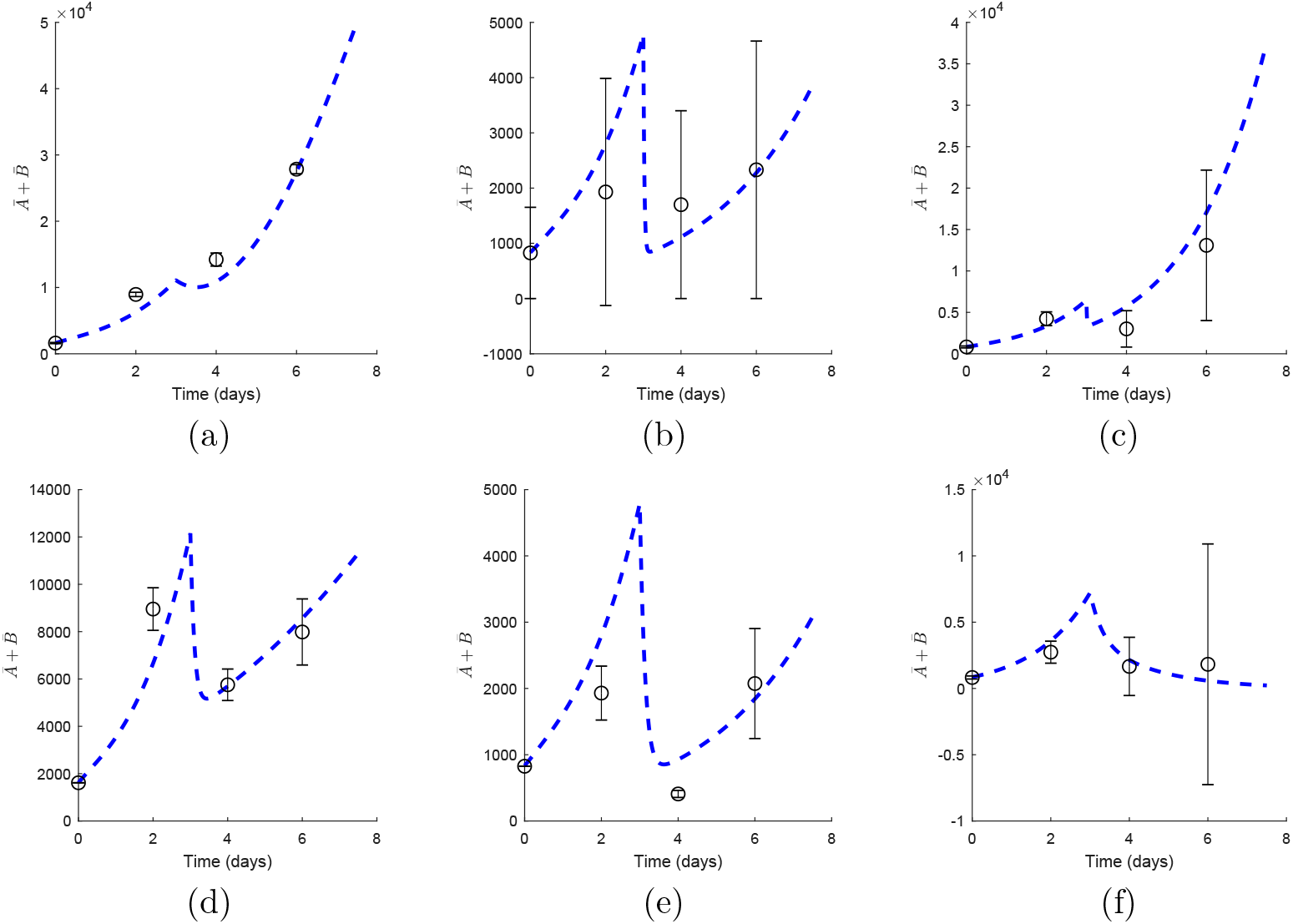
Fitting results of Equation (S2) to the WT, M1 and M2 data from [7] treated with afatinib and bortezomib. Figures (a), (b) and (c) show the WT, M1 and M2 data treated with afatinib, respectively. Figures (d), (e) and (f) show the WT, M1 and M2 data treated with bortezomib, respectively.

#### Treatment induced periodic environment

Most anti-cancer therapies include a recovery period following each treatment where the drug washes out and the patient recovers from the effects of treatment. Classical chemotherapy induces an approximately periodic tumour microenvironment with respect to the concentration of the chemotherapeutic agent, where each treatment cycle acts as the beginning of a new period. In what follows, we assume that the chemotherapeutic drug *C*(*t*) is administered periodically with a period of *T* days, and eliminated according to (S10) with elimination rate *k_elim_*. To facilitate the calculation of the reproduction number during treatment, we derive an estimate for the forced limit cycle in *C*(*t*) during periodic therapy.

Now, let the first dose be given at time *t* = *t_0_*, and assume that the administration time, *T_admin_* is negligible, so that each administration of therapy is given as an impulse at time *t_0_* + *n_T_*. Then, for 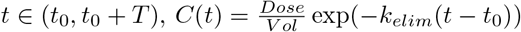. At time *t* = *T*, a second dose is given, so 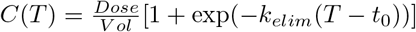, and the concentration of the drug decays, for *t* ∈ (*T, 2T*), according to *C*(*t*) = *C*(*T*) exp(—*k_elim_*(*t* — *T*)). Proceeding inductively, we see that, immediately after administering the *n* + 1st dose at *t* = *t_0_* + *nT*^+^,

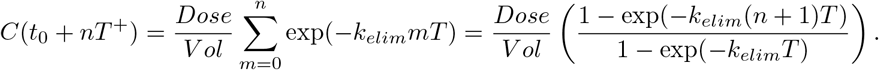

Then, for *t* ∈ (*t*_0_ + *nT, t*_0_ + (*n* + 1)*T*), *C*(*t*) = *C*(*nT^+^*) exp(—*k_elim_*(*t* — *NT*)). As the number of administrations, *n*, grows, the term exp(—*k_elim_*(*n* + 1)*T*) becomes increasingly small. Recalling that the half effect of the chemotherapeutic is given by

**Table S1.** The switching parameters for WT, M1, and M2 cell lines.

| Cell Type | Drug | $P_{BB}^{max}$ | $P_{BB}^*$ | $\sigma_A = \sigma_B$ (1/hour) | $d_A^{max}$ (1/hour) |
| --- | --- | --- | --- | --- | --- |
| WT | Docetaxel | 0.93 | 0.62 | $1 \times 10^{-2}$ | 0.7025 |
| WT | Afatinib | 0.95 | 0.81 | $1 \times 10^{-2}$ | 0.4828 |
| WT | Bortemozib | 0.95 | 0 | $1 \times 10^{-2}$ | 0.7993 |
| M1 | Docetaxel | 0.91 | 0 | $1 \times 10^{-2}$ | 1.145 |
| M1 | Afatinib | 0.92 | 0 | $1 \times 10^{-2}$ | 1.699 |
| M1 | Bortemozib | 92 | 0 | $1 \times 10^{-2}$ | 0.4491 |
| M2 | Docetaxel | 0.71 | 0.09 | $1 \times 10^{-2}$ | 0.1447 |

**Table S2.** The tumour growth parameters for the WT, M1 and M2 type cells.

| Cell Type | $r_A$ (1/hour) | $r_B$ (1/hour) | $d_A$ (1/hour) |
| --- | --- | --- | --- |
| WT | 0.4827 | 0.3498 | 0.4198 |
| M1 | 0.1646 | 0.0100 | 0.1163 |
| M2 | 0.1114 | 0.0857 | 0.0768 |

*C*_1/2_, we discard the influence of drug concentrations that are less than *ωC*_1/2_ for a given value of ω ≪ 1. Thus, after

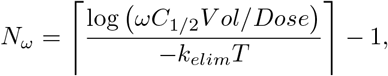

administrations, the error induced by discarding the exp(— *k_elim_*(*N_ω_* + 1)*T*) terms is

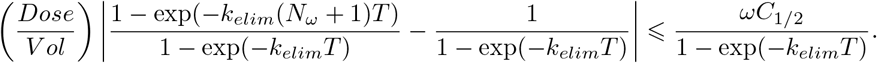

Accordingly, for a given value of *ω* and after the *N_ω_*-st administration of the chemotherapeutic, we consider the drug concentration in the tumour microenvironment to be in a periodically forced limit cycle given by

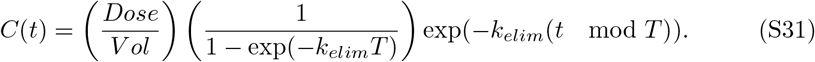

#### 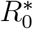 in the treated environment

Having derived an estimate for the chemotherapeutic concentration during metronomic therapy, we study the effect of this therapy on tumour growth. Once again, we are assuming that the drug tolerant population is not self-sustaining and has a negative intrinsic growth rate, λ*_B_* < 0, as in the NSCLC data considered. We assume that the chemotherapeutic has been administered at least *N_ω_* times so that the tumour microenvironment is roughly periodic and consider the age structured PDE

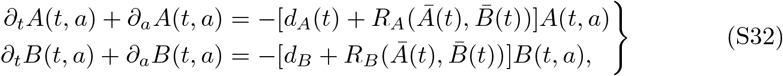

Once again, solving (S32) along the characteristic lines gives

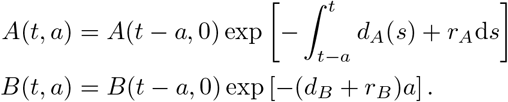

We thus obtain the corresponding renewal equations

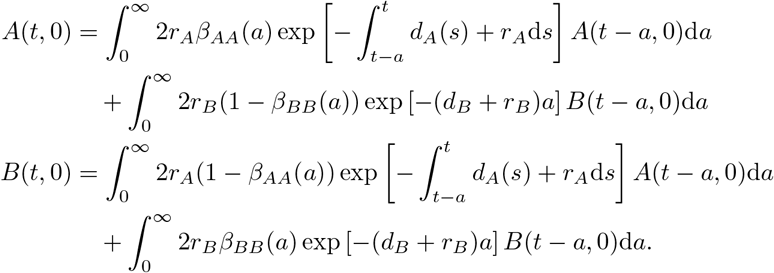

Taking the Laplace transform of these renewal equations gives the linear system

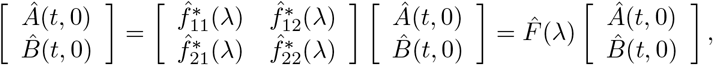

The treated NGO is therefore time dependent, due to the drug induced periodicity in the tumour microenvironment, and given by

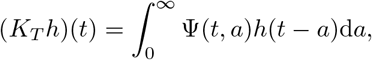

where, recalling that *β_ij_* (*a*) = 1 — *β_ij_*(*a*),

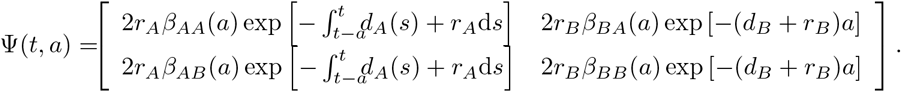

Now, it is important to note that periodic therapy does not immediately induce a periodic environment. However, as shown, for a large number of administrations, the error induced by assuming that *C*(*t*) is given by (S31) and can be made arbitrarily small. Moreover, we are interested in the asymptotic behaviour of the population of the tumour population, and therefore make the simplifying assumption that the drug concentration, and thus the environment, is effectively periodic.

The treated NGO *K_T_* acts on the space of *T*-periodic functions 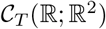. However, this is inconvenient for calculation purposes and we follow [47, 79] and pass the periodicity from the function *h* to the operator by defining

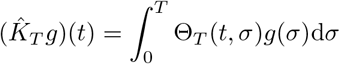

where 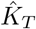 acts on 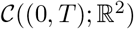, and Θ*_T_*(*t*,σ) is a periodic function defined by

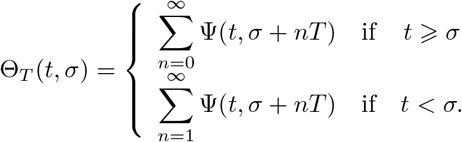

It follows that the spectral radius of *K_T_* equals that of 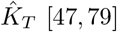 [47,79]. After interchanging the order of integration and summation, the treated basic reproduction number 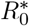 is given by the spectral radius of

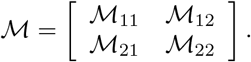

where

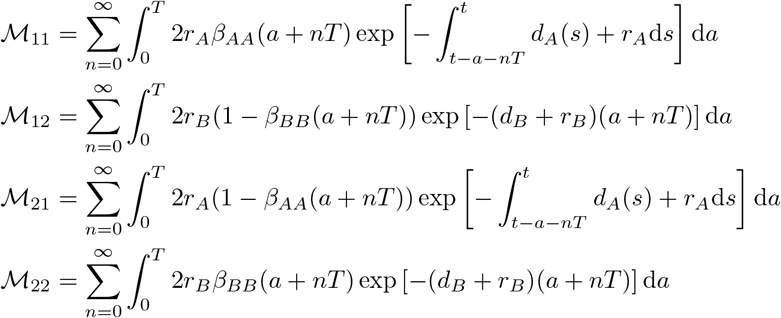

As the drug tolerant cells are immune to therapy, both *ℳ*_12_ and *ℳ*_22_ are constant in time, so these infinite series telescope and we immediately see *M*_12_ = *ℳ*_12_ and *M*_22_ = *ℳ*_22_, where *M*_*i*2_ is defined in the untreated NGO. It remains to calculate the effects of therapy on *ℳ*_11_ and *ℳ*_21_. Treatment increases the death rate of drug susceptible cells via

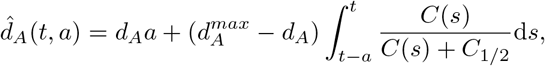

Using the estimate for *C*(*t*) during periodic therapy derived previously, it is possible to calculate 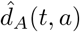 explicitly, and thus calculate 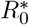. However, we recall that *R*_0_ = 1 is the threshold between disease growth and decay, and therefore the value of clinical interest. We now calculate a relationship between dose frequency, *T* and dose size to ensure that the treated reproduction number is below this threshold, so 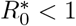. Accordingly, if

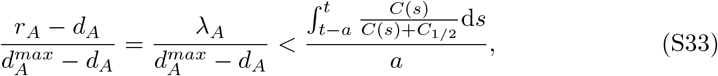

it follows that

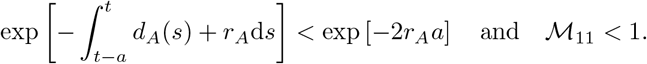

Then, we see that

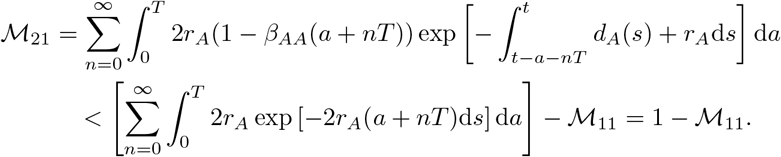

From the proof of Theorem S9, 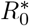 is a strictly increasing function of *ℳ*_21_, so it follows that

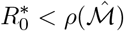

where 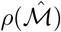 is the spectral radius of

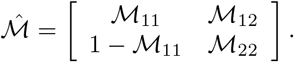

Using the fact that λ*_B_* < 0, the Gershgorin Circle Theorem gives

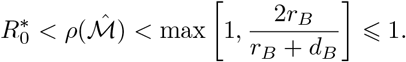

Thus, the condition (S33) is sufficient to ensure that 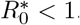. We now use the estimate for the periodic chemotherapeutic concentration derived in (S31) to derive a dose size to ensure that (S33) holds and, more importantly, that 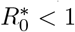. For notational simplicity, denote 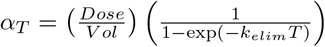 so

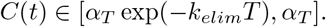

Now, let *C*\* be the solution of

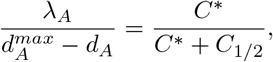

so

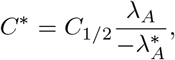

and a sufficient condition for (S33) to hold is

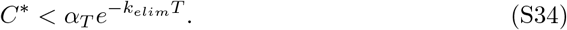

To see that (S34) is necessary for (S33) to hold, first note that if the required dose *C*\* satisfies *C*\* > *α_T_*, then it is not possible to administer a large enough dose to drive 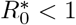 and (S33) cannot hold. Now, consider the case where that 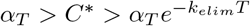. As *C*(*t*) attains the lower bound 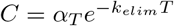 directly before the subsequent administration, it follows from the assumption 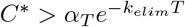 that there exists *t*\* such that *C*(*t*\*) < *C*\*. As *C*(*t*) is continuous and *α_T_* > *C*\*, there must be an *a*\* such that *C*(*t*\* — *a*\*) = *C*\*. Then, since *C*(*t*) is strictly decreasing between drug administrations, it follows that *C*(*s*) < *C*\* for *s* ∈ (*t*\* — *a*\*,*t*\*) and

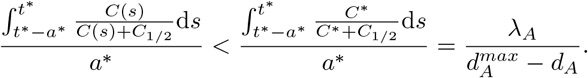

Consequently, 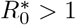 during the interval (*t*^*^ — *a*^*^, *t*^*^) for each administration period, and the tumour population may not decay. Thus, we conclude that the dose size must be chosen such that (S34) holds, which gives the threshold dose size

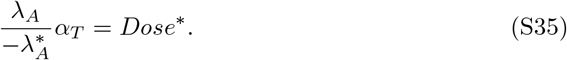

For a given chemotherapeutic agent, both *C*_1/2_ and *k_elim_* are fixed. Thus, the attending clinician can control the quantity 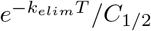 by varying treatment frequency and intensity.

#### Limiting cooperation of drug tolerant cells

It may be tempting to increase the amount of chemotherapeutic administered, and indeed (S35) appears to supports the usage of maximally tolerated dosing of anti-cancer drugs. However, this maximal dose size may allow for the competitive release of a drug-tolerant phenotype and the resulting resistance to therapy, which was not considered in the preceding analysis. Thus, therapy must be designed to balance the need to ensure 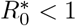 while guarding against the evolution of resistance. We incorporated the Allee effect and cooperation of drug tolerant cells in the mathematical model through the function *f_N_*(*θ*). Naively including cooperation, the fitness of the drug tolerant population is given by

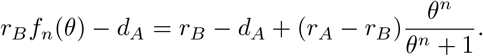

Thus, the threshold ratio *θ*\* must ensure *r_B_f_n_*(*θ*) — *d_A_* < 0. Using the definition of *f_n_*(*θ*) and re-arranging gives

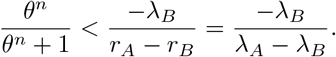

Then, a simple calculation gives the ratio θ^*^ defined in the main text.

#### Parametrization of chemotherapeutic pharmacokinetics

Docetaxel has an effective half life of roughly 86 hours [52], so we set *k_elim_* = log(2)/86/24 days^-1^ in our simple pharmacokinetic model, and an *in vitro* half-effect concentration of 4ng/mL [80], which is orders of magnitude less than the achievable plasma concentrations. Using the common dose size of 100mg/m^2^, and volume of distribution 74L/m^2^ [81], the ratio of Dose/*V ol* for half-maximum effect is roughly 10^4^. We used the 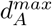 value calculated from the parameter fitting in the Methods to complete the pharmacodynamic model of docetaxel. To simulate the fixed therapy schedule, we set *T* = 7 days and fixed *C*_1/2_ = 0.5. For *V ol* = 74L/m^2^, the dose size during the fixed therapy schedule was calculated by satisfying Dose/*Vol*/*C*_1/2_ = 10^4^. As previously mentioned, it is this ratio and the value of 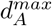 that determines the pharmacodynamics in our simple model.

To simulate the administration of afatinib, we set *t_1/2_* = 37 hours [82,83]. Afatinib is administered daily as an 40 mg oral capsule with *C_max_* = 25.2ng/mL [82,83]. The steady state concentration of daily afatinib is roughly 2.11 × *C_max_* [83]. As we do not update the model for *C*(*t*) to be specific for oral administration of drug, we can use the approximation for the steady state concentration of *C*(*t*) to calculate

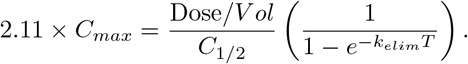

Then, once again we can calculate the ratio of Dose/*C*_1/2_, which along with the elimination constant *k_elim_* determines the pharmacokinetics and pharmacodynamics of afatinib.

To simulate the fixed periodic administration of bortezomib, we set *t*_1/2_ = 40 hours, and take *T* = 3 days to account for the minimum 72 hours between intravenous administrations [84]. Bortezomib has a large volume of distribution, between 498-1884 L/m^2^, so we take *V ol* = 850 L, while [84] report *C_max_* = 162 ng/mL following multiple I.V administrations. Thus, we once again use the approximation for the steady state concentration of *C*(*t*) to calculate the value of Dose.

#### Model informed therapy for other therapeutics

We implemented the model informed therapeutic strategy for afatinib and bortemozib in the WT and M1 cells and show the results in Figure S7. We note that the parameter estimates for the M2 population do not satisfy *r_B_* < *d_B_*, so the model informed therapy

**Table S3.** The effectiveness of model informed therapy when compared to periodic dosing over 150 days.

| Cell Type | Drug | Treatment Effectiveness |
| --- | --- | --- |
| WT | Afatinib | 0.9587 |
| WT | Bortemozib | 0.7640 |
| M1 | Afatinib | 0.4308 |
| M1 | Bortemozib | 0.4397 |

cannot be applied. For WT and M1 treated with afatinib and bortezomib, the increased effectiveness of the model informed therapy over 100 days of therapy is shown in Table S3.

We note that the model informed therapy for WT cells with afatinib does not lead to population extinction. For WT cells treated with afatinib, 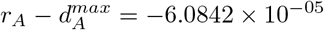. It follows that it must be the case that *C*(*t*) » *C*_1/2_ if (S34) is to be satisfied. Accordingly, *C*(*t*) decays too slowly between doses to inhibit the establishment of a drug tolerant population. Thus, the model informed therapy, while outperforming periodic dosing, does not drive tumour extinction.

**Fig S7.**
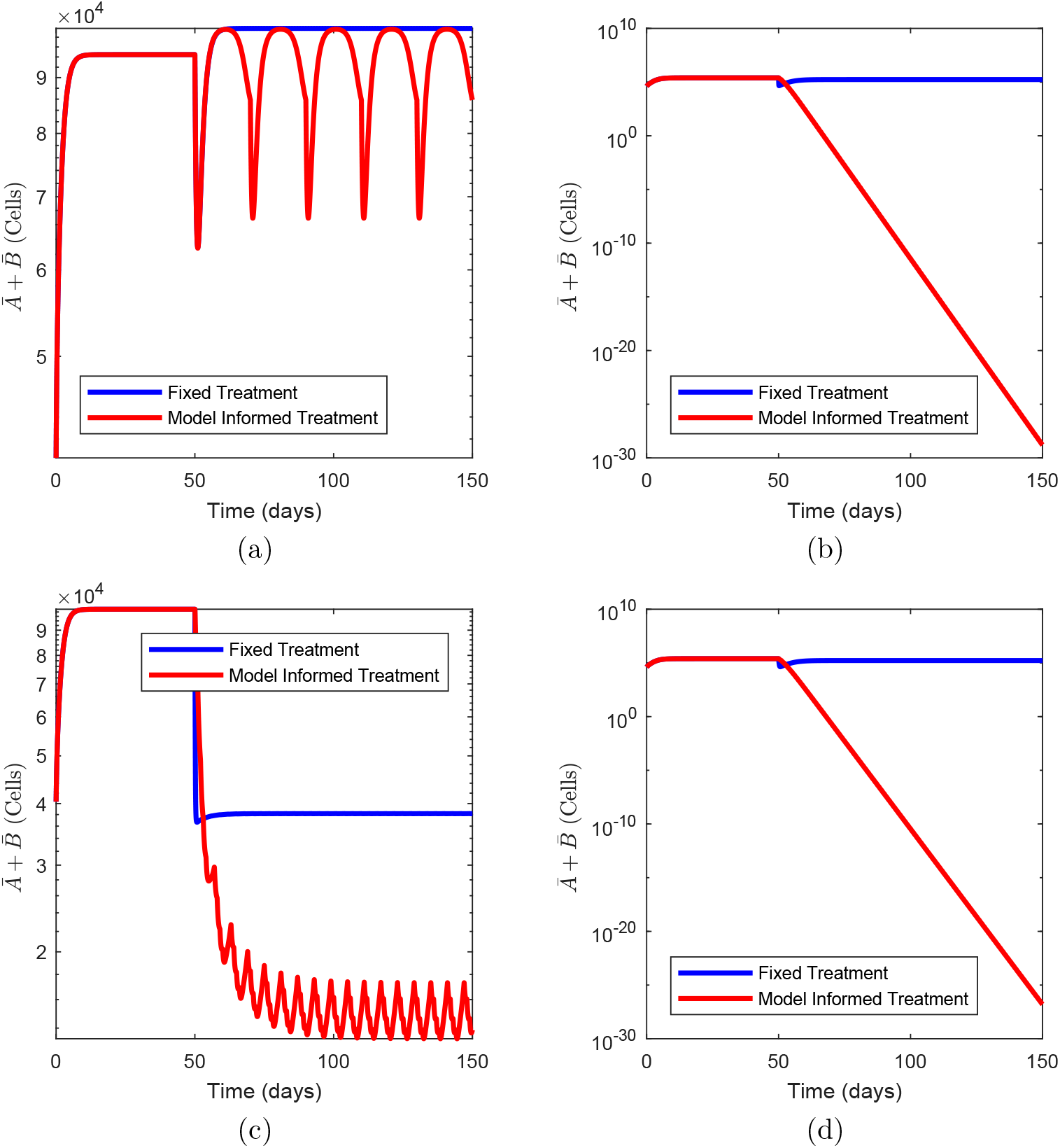
Comparison of periodic therapy in blue and model informed therapy in red for afatinib and bortezomib. Figures (a) and (b) are the WT and M1 cells treated with afatinib, respectively. Figures (c) and (d) are the WT and M1 cells treated with bortemozib, respectively.

## References

1. Greaves M. Evolutionary Determinants of Cancer. Cancer Discovery. 2015;5(8):806–820. doi:10.1158/2159-8290.CD-15-0439.

2. Korolev KS, Xavier JB, Gore J. Turning ecology and evolution against cancer. Nature Reviews Cancer. 2014;14(5):371–380. doi:10.1038/nrc3712.

3. Dagogo-Jack I, Shaw AT. Tumour heterogeneity and resistance to cancer therapies. Nature Reviews Clinical Oncology. 2018;15(2):81–94. doi:10.1038/nrclinonc.2017.166.

4. Marusyk A, Polyak K. Tumor heterogeneity: Causes and consequences. Biochimica et Biophysica Acta (BBA) - Reviews on Cancer. 2010;1805(1):105–117. doi:10.1016/j.bbcan.2009.11.002.

5. Gatenby RA, Silva AS, Gillies RJ, Frieden BR. Adaptive Therapy. Cancer Research. 2009;69(11):4894–4903. doi:10.1158/0008-5472.CAN-08-3658.

6. Altrock PM, Liu LL, Michor F. The mathematics of cancer: integrating quantitative models. Nature Reviews Cancer. 2015;15(12):730–745. doi:10.1038/nrc4029.

7. Craig M, Kaveh K, Woosley A, Brown AS, Goldman D, Eton E, et al. Cooperative adaptation to therapy (CAT) confers resistance in heterogeneous non-small cell lung cancer. PLOS Computational Biology. 2019;15(8):e1007278. doi:10.1371/journal.pcbi.1007278.

8. Ramirez M, Rajaram S, Steininger RJ, Osipchuk D, Roth MA, Morinishi LS, et al. Diverse drug-resistance mechanisms can emerge from drug-tolerant cancer persister cells. Nature Communications. 2016;7(1):10690. doi:10.1038/ncomms10690.

9. Holohan C, Van Schaeybroeck S, Longley DB, Johnston PG. Cancer drug resistance: an evolving paradigm. Nature Reviews Cancer. 2013;13(10):714–726. doi:10.1038/nrc3599.

10. Horvath D, Brutovsky B. Toward understanding of the role of reversibility of phenotypic switching in the evolution of resistance to therapy. Physics Letters A. 2018;382(24):1586–1600. doi:10.1016/j.physleta.2018.03.052.

11. Chisholm RH, Lorenzi T, Clairambault J. Cell population heterogeneity and evolution towards drug resistance in cancer: Biological and mathematical assessment, theoretical treatment optimisation. Biochimica et Biophysica Acta (BBA) - General Subjects. 2016;1860(11):2627–2645. doi:10.1016/j.bbagen.2016.06.009.

12. Gallaher J, Anderson ARA. Evolution of intratumoral phenotypic heterogeneity: The role of trait inheritance. Interface Focus. 2013;3(4). doi:10.1098/rsfs.2013.0016.

13. Nichol D, Robertson-Tessi M, Jeavons P, Anderson ARA. Stochasticity in the Genotype-Phenotype Map: Implications for the Robustness and Persistence of Bet-Hedging. Genetics. 2016;204(4):1523–1539. doi:10.1534/genetics.116.193474.

14. Jolly MK, Kulkarni P, Weninger K, Orban J, Levine H. Phenotypic Plasticity, Bet-Hedging, and Androgen Independence in Prostate Cancer: Role of Non-Genetic Heterogeneity. Frontiers in Oncology. 2018;8(MAR):1–12. doi:10.3389/fonc.2018.00050.

15. Sharma SV, Lee DY, Li B, Quinlan MP, Takahashi F, Maheswaran S, et al. A Chromatin-Mediated Reversible Drug-Tolerant State in Cancer Cell Subpopulations. Cell. 2010;141(1):69–80. doi:10.1016/j.cell.2010.02.027.

16. Goldman A, Majumder B, Dhawan A, Ravi S, Goldman D, Kohandel M, et al. Temporally sequenced anticancer drugs overcome adaptive resistance by targeting a vulnerable chemotherapy-induced phenotypic transition. Nature Communications. 2015;6(1):6139. doi:10.1038/ncomms7139.

17. Pigliucci M. Phenotypic Plasticity: Beyond Nature and Nurture. Baltimore: Johns Hopkins University Press; 2001.

18. King OD, Masel J. The evolution of bet-hedging adaptations to rare scenarios. Theoretical population biology. 2007;72(4):560–75. doi:10.1016/j.tpb.2007.08.006.

19. Harms A, Maisonneuve E, Gerdes K. Mechanisms of bacterial persistence during stress and antibiotic exposure. Science. 2016;354(6318):aaf4268. doi:10.1126/science.aaf4268.

20. Lewis K. Persister cells, dormancy and infectious disease. Nature Reviews Microbiology. 2007;5(1):48–56. doi:10.1038/nrmicro1557.

21. Muller J, Hense BA, Fuchs TM, Utz M, Potzsche C. Bet-hedging in stochastically switching environments. Journal of Theoretical Biology. 2013;336:144–157. doi:10.1016/j.jtbi.2013.07.017.

22. Zhang J, Cunningham JJ, Brown JS, Gatenby RA. Integrating evolutionary dynamics into treatment of metastatic castrate-resistant prostate cancer. Nature Communications. 2017;8(1):1816. doi:10.1038/s41467-017-01968-5.

23. Bacevic K, Noble R, Soffar A, Wael Ammar O, Boszonyik B, Prieto S, et al. Spatial competition constrains resistance to targeted cancer therapy. Nature Communications. 2017;8(1):1995. doi:10.1038/s41467-017-01516-1.

24. West J, You L, Zhang J, Gatenby RA, Brown JS, Newton PK, et al. Towards multi-drug adaptive therapy. Cancer Research. 2020; p. canres.2669.2019. doi:10.1158/0008-5472.CAN-19-2669.

25. Pigliucci M. Genotype-phenotype mapping and the end of the ‘genes as blueprint’ metaphor. Philosophical Transactions of the Royal Society B: Biological Sciences. 2010;365(1540):557–566. doi:10.1098/rstb.2009.0241.

26. Nichol D, Robertson-Tessi M, Anderson ARA, Jeavons P. Model genotype-phenotype mappings and the algorithmic structure of evolution. Journal of The Royal Society Interface. 2019;16(160):20190332. doi:10.1098/rsif.2019.0332.

27. Ardaseva A, Gatenby RA, Anderson ARA, Byrne HM, Maini PK, Lorenzi T. Evolutionary dynamics of competing phenotype-structured populations in periodically fluctuating environments. Journal of Mathematical Biology. 2019;doi:10.1007/s00285-019-01441-5.

28. Lorenzi T, Chisholm RH, Clairambault J. Tracking the evolution of cancer cell populations through the mathematical lens of phenotype-structured equations. Biology Direct. 2016;11(1):1–17. doi:10.1186/s13062-016-0143-4.

29. Busse JE, Gwiazda P, Marciniak-Czochra A. Mass concentration in a nonlocal model of clonal selection. Journal of Mathematical Biology. 2016;73(4):1001–1033. doi:10.1007/s00285-016-0979-3.

30. Chisholm RH, Lorenzi T, Lorz A, Larsen AK, De Almeida LN, Escargueil A, et al. Emergence of Drug Tolerance in Cancer Cell Populations: An Evolutionary Outcome of Selection, Nongenetic Instability, and Stress-Induced Adaptation. Cancer Research. 2015;75(6):930–939. doi:10.1158/0008-5472.CAN-14-2103.

31. Yang HW, Chung M, Kudo T, Meyer T. Competing memories of mitogen and p53 signalling control cell-cycle entry. Nature. 2017;549(7672):404–408. doi:10.1038/nature23880.

32. Cardelli L, Csikász-Nagy A. The Cell Cycle Switch Computes Approximate Majority. Scientific Reports. 2012;2(1):656. doi:10.1038/srep00656.

33. Cardelli L. Morphisms of reaction networks that couple structure to function. BMC Systems Biology. 2014;8(1):84. doi:10.1186/1752-0509-8-84.

34. Keren I, Kaldalu N, Spoering A, Wang Y, Lewis K. Persister cells and tolerance to antimicrobials. FEMS Microbiology Letters. 2004;230(1):13–18. doi:10.1016/S0378-1097(03)00856-5.

35. Balaban NQ, Merrin J, Chait R, Kowalik L, Leibler S. Bacterial persistence as a phenotypic switch; Supplemental Materials. Science. 2004;305(5690):1622–1625.

36. Shaffer SM, Dunagin MC, Torborg SR, Torre EA, Emert B, Krepler C, et al. Rare cell variability and drug-induced reprogramming as a mode of cancer drug resistance. Nature. 2017;546(7658):431–435. doi:10.1038/nature22794.

37. Pisco AO, Huang S. Non-genetic cancer cell plasticity and therapy-induced stemness in tumour relapse: ‘What does not kill me strengthens me’. British Journal of Cancer. 2015;112(11):1725–1732. doi:10.1038/bjc.2015.146.

38. Gallie J, Libby E, Bertels F, Remigi P, Jendresen CB, Ferguson GC, et al. Bistability in a Metabolic Network Underpins the De Novo Evolution of Colony Switching in Pseudomonas fluorescens. PLoS Biology. 2015;13(3):1–28. doi:10.1371/journal.pbio.1002109.

39. Gravenmier CA, Siddique M, Gatenby RA. Adaptation to Stochastic Temporal Variations in Intratumoral Blood Flow: The Warburg Effect as a Bet Hedging Strategy. Bulletin of Mathematical Biology. 2018;80(5):954–970. doi:10.1007/s11538-017-0261-x.

40. Dingli D, Chalub FACC, Santos FC, Van Segbroeck S, Pacheco JM. Cancer phenotype as the outcome of an evolutionary game between normal and malignant cells. British Journal of Cancer. 2009;101(7):1130–1136. doi:10.1038/sj.bjc.6605288.

41. Ross-Gillespie A, Gardner A, Buckling A, West SA, Griffin AS. Density Dependence and CooperationL Theory and a Test with Bacteria. Evolution. 2009;63(9):2315–2325. doi:10.1111/j.1558-5646.2009.00723.x.

42. Kimmel GJ, Gerlee P, Brown JS, Altrock PM. Neighborhood size-effects shape growing population dynamics in evolutionary public goods games. Communications Biology. 2019;2(1):53. doi:10.1038/s42003-019-0299-4.

43. Archetti M, Pienta KJ. Cooperation among cancer cells: applying game theory to cancer. Nature Reviews Cancer. 2019;19(2):110–117. doi:10.1038/s41568-018-0083-7.

44. Proenca AM, Rang CU, Buetz C, Shi C, Chao L. Age structure landscapes emerge from the equilibrium between aging and rejuvenation in bacterial populations. Nature Communications. 2018;9(1):3722. doi:10.1038/s41467-018-06154-9.

45. Govers SK, Adam A, Blockeel H, Aertsen A. Rapid phenotypic individualization of bacterial sister cells. Scientific Reports. 2017;7(1):8473. doi:10.1038/s41598-017-08660-0.

46. Diekmann O, Heesterbeek JAP, Metz JAJ. On the definition and the computation of the basic reproduction ratio R 0 in models for infectious diseases in heterogeneous populations. Journal of Mathematical Biology. 1990;28(4):365–382. doi:10.1007/BF00178324.

47. Inaba H. The Malthusian parameter and $R_0$ for heterogeneous populations in periodic environments. Mathematical Biosciences and Engineering. 2012;9(2):313–346. doi:10.3934/mbe.2012.9.313.

48. Inaba H. On a new perspective of the basic reproduction number in heterogeneous environments. Journal of Mathematical Biology. 2012;65(2):309–348. doi:10.1007/s00285-011-0463-z.

49. British Colombia Cancer Agency. The Cancer Drug Manual. Vancouver, Canada: Provincial Health Services Authority; 2019.

50. Siegel RL, Miller KD, Jemal A. Cancer statistics, 2018. CA Cancer J Clin. 2018;68(1):7–30. doi:10.3322/caac.21442.

51. Rotow J, Bivona TG. Understanding and targeting resistance mechanisms in NSCLC. Nat Rev Cancer. 2017;17(11):637–658. doi:10.1038/nrc.2017.84.

52. Baker SD, Zhao M, Lee CKK, Verweij J, Zabelina Y, Brahmer JR, et al. Comparative Pharmacokinetics of Weekly and Every-Three-Weeks Docetaxel. Clinical Cancer Research. 2004;10(6):1976–1983. doi:10.1158/1078-0432.CCR-0842-03.

53. Min M, Spencer SL. Spontaneously slow-cycling subpopulations of human cells originate from activation of stress-response pathways. PLOS Biol. 2019;17(3):e3000178. doi:10.1371/journal.pbio.3000178.

54. Shaffer SM, Emert BL, Reyes Hueros RA, Cote C, Harmange G, Schaff DL, et al. Memory Sequencing Reveals Heritable Single-Cell Gene Expression Programs Associated with Distinct Cellular Behaviors. Cell. 2020;182(4):947–959.e17. doi:10.1016/j.cell.2020.07.003.

55. Chakrabarti S, Paek AL, Reyes J, Lasick KA, Lahav G, Michor F. Hidden heterogeneity and circadian-controlled cell fate inferred from single cell lineages. Nat Commun. 2018;9(1):5372. doi:10.1038/s41467-018-07788-5.

56. Gallaher JA, Enriquez-Navas PM, Luddy KA, Gatenby RA, Anderson ARA. Spatial Heterogeneity and Evolutionary Dynamics Modulate Time to Recurrence in Continuous and Adaptive Cancer Therapies. Cancer Research. 2018;78(8):2127–2139. doi:10.1158/0008-5472.CAN-17-2649.

57. Strobl MAR, Gallaher J, West J, Robertson-Tessi M, Maini PK, Anderson ARA. Spatial structure impacts adaptive therapy by shaping intra-tumoral competition. bioRxiv. 2020;(11.03.365163):1–31. doi:10.1101/2020.11.03.365163.

58. Robertson-Tessi M, Gillies RJ, Gatenby RA, Anderson ARA. Impact of metabolic heterogeneity on tumor growth, invasion, and treatment outcomes. Cancer Res. 2015;75(8):1567–1579. doi:10.1158/0008-5472.CAN-14-1428.

59. Karolak A, Rejniak KA. Micropharmacology: An In Silico Approach for Assessing Drug Efficacy Within a Tumor Tissue. Bull Math Biol. 2019;81(9):3623–3641. doi:10.1007/s11538-018-0402-x.

60. Nicholson MD, Antal T. Competing evolutionary paths in growing populations with applications to multidrug resistance. PLoS Comput Biol. 2019;15(4):e1006866. doi:10.1371/journal.pcbi.1006866.

61. Kaznatcheev A, Peacock J, Basanta D, Marusyk A, Scott JG. Fibroblasts and alectinib switch the evolutionary games played by non-small cell lung cancer. Nat Ecol Evol. 2019;3(3):450–456. doi:10.1038/s41559-018-0768-z.

62. Chen S, Xue Y, Wu X, Le C, Bhutkar A, Bell EL, et al. Global microRNA depletion suppresses tumor angiogenesis. Genes Dev. 2014;28(10):1054–1067. doi:10.1101/gad.239681.114.

63. Perthame B. Transport Equations in Biology. Frontiers in Mathematics. Basel: Birkhäuser Basel; 2007. Available from: http://www.amazon.com/exec/obidos/redirect?tag=citeulike07-20{&}path=ASIN/3764378417http://link.springer.com/10.1007/978-3-7643-7842-4.

64. MATLAB. R2017a. Natick, Massachusetts: The MathWorks Inc.; 2017.

65. Cassidy T, Craig M, Humphries AR. Equivalences between age structured models and state dependent distributed delay differential equations. Mathematical Biosciences and Engineering. 2019;16(5):5419–5450. doi:10.3934/mbe.2019270.

66. Rundnicki R, Mackey MC. Asymptotic Similarity and Malthusian Growth in Autonomous and Nonautonomous Populations. Journal of Mathematical Analysis and Applications. 1994;187(2):548–566. doi:10.1006/jmaa.1994.1374.

67. Billy F, Clairambaultt J, Fercoq O, Gaubertt S, Lepoutre T, Ouillon T, et al. Synchronisation and control of proliferation in cycling cell population models with age structure. Mathematics and Computers in Simulation. 2014;96:66–94. doi:10.1016/j.matcom.2012.03.005.

68. Cassidy T, Humphries AR. A mathematical model of viral oncology as an immuno-oncology instigator. Mathematical Medicine and Biology: A Journal of the IMA. 2019;To appear. doi:10.1093/imammb/dqz008.

69. Metz JAJ, Diekmann O, editors. The Dynamics of Physiologically Structured Populations. vol. 68 of Lecture Notes in Biomathematics. 3rd ed. Berlin, Heidelberg: Springer Berlin Heidelberg; 1986. Available from: http://link.springer.com/10.1007/978-3-662-13159-6.

70. Greaves M, Maley CC. Clonal evolution in cancer. Nature. 2012;481(7381):306–313. doi:10.1038/nature10762.

71. Ben-Jacob E, S Coffey D, Levine H. Bacterial survival strategies suggest rethinking cancer cooperativity. Trends in Microbiology. 2012;20(9):403–410. doi:10.1016/j.tim.2012.06.001.

72. Gillies RJ, Brown JS, Anderson ARA, Gatenby RA. Eco-evolutionary causes and consequences of temporal changes in intratumoural blood flow. Nature Reviews Cancer. 2018;18(9):576–585. doi:10.1038/s41568-018-0030-7.

73. Andersson DI, Hughes D. Antibiotic resistance and its cost: is it possible to reverse resistance? Nature Reviews Microbiology. 2010;8(4):260–271. doi:10.1038/nrmicro2319.

74. Diekmann O, Gyllenberg M, Metz JAJ. Finite dimensional state representation of linear and nonlinear delay systems. Journal of Dynamics and Differential Equations. 2018;30(4):1439–1467. doi:10.1007/s10884-017-9611-5.

75. Concepción-Acevedo J, Weiss HN, Chaudhry WN, Levin BR. Malthusian Parameters as Estimators of the Fitness of Microbes: A Cautionary Tale about the Low Side of High Throughput. PLOS ONE. 2015;10(6):e0126915. doi:10.1371/journal.pone.0126915.

76. Inaba H, Nishiura H. The state-reproduction number for a multistate class age structured epidemic system and its application to the asymptomatic transmission model. Mathematical Biosciences. 2008;216(1):77–89. doi:10.1016/j.mbs.2008.08.005.

77. Inaba H. On the definition and the computation of the type-reproduction number T for structured populations in heterogeneous environments. Journal of Mathematical Biology. 2013;66(4-5):1065–1097. doi:10.1007/s00285-012-0522-0.

78. Dingli D, Offord C, Myers R, Peng KW, Carr TW, Josic K, et al. Dynamics of multiple myeloma tumor therapy with a recombinant measles virus. Cancer Gene Therapy. 2009;16(12):873–882. doi:10.1038/cgt.2009.40.

79. Bacaёr N, Ait Dads EH. Genealogy with seasonality, the basic reproduction number, and the influenza pandemic. Journal of Mathematical Biology. 2011;62(5):741–762. doi:10.1007/s00285-010-0354-8.

80. Bissery MC. Preclinical pharmacology of docetaxel. European Journal of Cancer. 1995;31(1004):S1–S6. doi:10.1016/0959-8049(95)00357-O.

81. Clarke SJ, Rivory LP. Clinical Pharmacokinetics of Docetaxel. Clinical Pharmacokinetics. 1999;36(2):99–114. doi:10.2165/00003088-199936020-00002.

82. Wind S, Schnell D, Ebner T, Freiwald M, Stopfer P. Clinical Pharmacokinetics and Pharmacodynamics of Afatinib. Clinical Pharmacokinetics. 2017;56(3):235–250. doi:10.1007/s40262-016-0440-1.

83. Agency EM. Giotrif: summary of product characteristics; 2018. Available from: https://www.ema.europa.eu/en/documents/product-information/giotrif-epar-product-information{_}en.pdf.

84. Tan CRC, Abdul-Majeed S, Cael B, Barta SK. Clinical Pharmacokinetics and Pharmacodynamics of Bortezomib. Clinical Pharmacokinetics. 2019;58(2):157–168. doi:10.1007/s40262-018-0679-9.

